# Oncodrive3D: Fast and accurate detection of structural clusters of somatic mutations under positive selection

**DOI:** 10.1101/2025.01.11.632354

**Authors:** Stefano Pellegrini, Olivia Dove Estrella, Ferran Muiños, Nuria Lopez-Bigas, Abel Gonzalez-Perez

## Abstract

Identifying the genes capable of driving tumorigenesis in different tissues is one of the central goals of cancer genomics. Computational methods that exploit different signals of positive selection in the pattern of mutations observed in genes across tumors have been developed to this end. One such signal of positive selection is clustering of mutations in areas of the three-dimensional (3D) structure of the protein above the expectation under neutrality. Methods that exploit this signal have been hindered by the paucity in proteins with experimentally solved 3D structures covering their entire sequence. Here, we present Oncodrive3D, a computational method that by exploiting AlphaFold 2 structural models extends the identification of proteins with significant mutational 3D clusters to the entire human proteome. Oncodrive3D shows sensitivity and specificity on par with state-of-the-art cancer driver gene identification methods exploiting mutational clustering, and clearly outperforms them in computational efficiency. We demonstrate, through several examples, how significant mutational 3D clusters identified by Oncodrive3D in different known or potential cancer driver genes can reveal details about the mechanism of tumorigenesis in different cancer types and in clonal hematopoiesis.

## Introduction

The identification of mutational cancer driver genes –those capable of driving tumorigenesis upon point mutations and indels– is one of the most important research goals of cancer genomics. The availability of somatic point mutations (mutations, for short) detected across cohorts of tumors, driven by the revolution of next generation sequencing technologies in the past two decades, led to the development of dozens of computational methods aimed at this goal (1–12). All these methods rely on the comparison of certain features of the mutations observed across tumors with the same features expected under neutral mutagenesis. Among the features exploited we can count the recurrence of mutations with a given consequence type, the average functional impact of mutations, and the distribution of mutations along the sequence or the three-dimensional structure of the protein. Significant differences between the observation and the expectation under neutrality for these features are often referred to as signals of positive selection. The values of these features in different cancer genes also provide information about the underlying mechanisms of tumorigenesis. For example, while loss-of-function genes tend to accumulate more truncating mutations than expected along their sequence, gain-of-function genes usually accumulate missense mutations at specific positions (13, 14).

We and others have shown that the combination of computational methods exploiting different signals of positive selection has brought within reach the complete compendium of mutational cancer driver genes across tumor types (15–18). As tumor mutation datasets available in the public domain become larger and comprise more malignancies (15, 19–21), the completion of the compendium requires methods of detection of signals of positive selection that fulfill at least three criteria: they must build accurate models of background mutagenesis to guarantee the reliability of their measurements of positive selection, be applicable to all mutated genes in the cohort and possess computational efficiency that guarantees scalability. While for most signals of positive selection methods fulfilling these three criteria have been developed –e.g., recurrence above expectation by dNdScv (3), linear clustering by OncodriveCLUSTL (11), functional impact bias of mutations by OncodriveFML (6)– this has proven harder for methods that specifically exploit the clustering of mutations within the three-dimensional structure of proteins (10, 22, 23). The paucity in experimentally resolved high-quality structure for most proteins in the human proteome has limited the possibility to detect three-dimensional (3D) mutational clusters to only a few hundreds of genes (24). Moreover, mutational 3D cluster methods can be computationally very expensive when compared with methods exploiting other signals of positive selection, if they map mutations to all structure fragments covering the sequence of a gene. Finally, their calculations of neutral mutagenesis do not generally incorporate the observed trinucleotide change frequencies, which are one of the most important determinants of the mutation rate across genes.

A second important output from the calculation of genes exhibiting significant clusters of mutations in their tertiary structure is the description (location, recurrence across tumor types, etc) of such clusters. They provide crucial information about the underlying mechanisms of tumorigenesis and support the identification of driver mutations (25). Previous efforts to identify clusters of mutations in cancer genes have focused mostly on mutational hotspots, which are easier to identify (26, 27).

To solve these hurdles, we have developed Oncodrive3D (https://github.com/bbglab/oncodrive3d), a fast and accurate novel 3D clustering algorithm for the discovery of cancer driver genes. Oncodrive3D leverages AlphaFold 2 (AlphaFold, for short) (28) structure predictions and predicted aligned error (PAE) to construct contact probability maps and to identify mutational clusters in the three-dimensional structure of proteins across all mutated genes in a cohort of tumors. To do that, it employs the trinucleotide mutational profiles observed across tumors to simulate neutral mutagenesis, and rank-based statistics to compute empirical p-values to detect residues exhibiting an accumulation of mutations in their 3D structural vicinity that is significantly beyond that expected under neutrality. Oncodrive3D then infers clumps of residues with significant accumulation of mutations that are close in the protein three-dimensional structure. In a study across 28,067 tumors of 83 cancer types, Oncodrive3D performs on par with a state-of-the-art 3D clustering-based driver discovery method, extending the discovery to the entire human proteome, and using 33 times less CPU-days. It complements other driver discovery methods based on different signals of positive selection, and is thus able to support the systematic effort to complete the compendium of mutational cancer driver genes. Furthermore, Oncodrive3D provides detailed annotations of mutational clusters in the three-dimensional structure of cancer proteins that appear under positive selection.

## Materials and Methods

### Somatic mutation data

Somatic mutations of 215 cohorts of tumors sequenced at the whole-genome and whole-exome level as part of different cancer genomics projects (including 32 by The Cancer Genome Atlas, TCGA) were obtained from the intOGen platform (17) (www.intogen.org). Blood somatic mutations identified across individuals in three separate cohorts (18, 21, 29–31) were obtained from the same source (www.intogen.org/ch). Mutations from hypermutated samples, identified as outliers in their respective cohorts as described in (17), had already been discarded by the intOGen pipeline. The pipeline had also mapped the mutations to the MANE Select transcripts (32) of each gene using ENSEMBL VEP v.111 (33). The number of samples and somatic mutations of all these cohorts are described in detail in Supplementary Table 1.

### AlphaFold models and related information

PDB files of AlphaFold protein structure models were obtained from the AlphaFold Protein Structure Database (28, 34) (https://alphafold.ebi.ac.uk) (AlphaFold 2 v.4). We downloaded structure models corresponding to MANE Select transcripts whenever possible (17,334 structures). For cases where models of corresponding MANE Select transcripts were not available, we complemented our dataset with available structures from the Human proteome in the AlphaFold Database (reference proteome UP000005640, which includes 23,391 structures) to ensure comprehensive coverage of the human proteome. By merging the two sets of structures, the total number of structural models of human proteins reached 24,349. Structural models of fragments of big proteins were combined using their overlapping sequences into full-size models, following methods previously described by others (35). We also obtained the Predicted Alignment Error (PAE) and the predicted local distance difference test (pLDDT) metrics from the AlphaFold Protein Structure database (https://alphafold.ebi.ac.uk). Solvent accessibility at the residue level and secondary structure information were extracted from the PDB structures using PDB_Tool (https://github.com/realbigws/PDB_Tool). The predicted stability change for each observed mutation was obtained from RaSP (36) (https://sid.erda.dk/cgi-sid/ls.py?share_id=fFPJWflLeE). The information on protein functional domains was downloaded from Ensembl using the BioMart service (37) (https://www.ensembl.org/info/data/biomart), while information on degron motifs was obtained from DegronMD (38) (https://bioinfo.uth.edu/degronmd).

The DNA sequence corresponding to each protein structure was obtained as follows. For genes mapped to an AlphaFold model structure corresponding to a UniProt ID with available coordinate information, we used the EMBL-EBI Protein API (39) (https://www.ebi.ac.uk/proteins/api/doc/) to retrieve the coding region coordinates. These coordinates were then used with bgreference (https://bitbucket.org/bgframework/bgreference/) to extract the DNA sequence, including the flanking regions at splicing sites required for the trinucleotide context. For genes not fulfilling the above criteria but mapped to an AlphaFold model structure with their corresponding Ensembl transcript ID, the DNA sequence was obtained using the Ensembl Get Sequence REST API (37) (https://rest.ensembl.org/). For any other genes the DNA sequence was obtained using the EMBL-EBI EMBOSS Backtranseq API (https://www.ebi.ac.uk/jdispatcher/st/emboss_backtranseq) (40).

### Other datasets used in downstream analyses

We downloaded manually curated bona fide cancer driver genes lists from the Cancer Gene Census v.99 (41, 42) (CGC; https://cancer.sanger.ac.uk/census) and OncoKB (available version at 12 July 2024) (43) (https://www.oncokb.org/cancer-genes). The former was used as a list of known cancer genes to benchmark the sensitivity of Oncodrive3D (see below), while the latter was used as annotations in several figures describing the results of the application of Oncodrive3D to different cohorts of tumors. We obtained a list of genes that are deemed highly unlikely to be involved in tumorigenesis across tissues, but which are frequently falsely identified by driver discovery methods due to their length or high mutation rate –for example, because they are not expressed in the tissue in question (15)– from intOGen (17) (https://www.intogen.org/faq). We called this the list of “Fishy” genes and used it to benchmark the specificity of Oncodrive3D. We actually produce different lists of Fishy genes per tissue, as the set of unexpressed genes vary between tissues. The classification of *bona fide* cancer genes into oncogenes and tumor suppressors was obtained from OncoKB (43), and TP53 was manually changed from ambiguous to tumor suppressor.

### Calculation of contact maps

Intuitively, two residues may be considered to be in contact with each other if the distance between their alpha carbons is below a given threshold. We reasoned that the confidence in the distance between any two residues in the protein structure model built by AlphaFold would be affected by their Predicted Alignment Error (PAE). Specifically, the PAE for the residue pair (C, R) represents the expected error at predicting the position of C if the true and predicted structures were aligned on residue R (28). Thus, higher PAE values for a pair of residues indicate greater uncertainty regarding their proximity.

To account for this uncertainty, we used the PAE to quantify the contact probability between a center residue C and any other residue R. For residue C we defined a spherical volume 𝑆_C_ centered at its alpha carbon with radius 10 Å, which gives a convenient estimate of the typical spatial extent of a residue’s side chain and therefore represents a valid measure to assess physical proximity. For residue *R*, we defined a spherical volume 𝑆_R_ centered at its alpha carbon with radius equal to the PAE value of the residue pair (R,C). We then calculated the intersection volume between 𝑆_R_ and 𝑆_C_ as the sum of the volumes of the respective spherical caps delimited by the intersection plane. Assuming that the position of R is uniformly distributed with support in the volume 𝑆_R_, the probability that C and R are in contact (closer that 10 Å) is given by 𝑣𝑜𝑙(𝑆_𝑅_ ∩ 𝑆_𝐶_)/𝑣𝑜𝑙(𝑆_𝑅_). This approach provides a contact probability that accounts for structural uncertainty introduced by the PAE, rather than relying solely on a fixed distance threshold. Finally, we defined residues as being in contact if their contact probability exceeded 0.5.

### Mutational profile

The 96-wide profile of trinucleotide changes, representing the propensity of each possible reference nucleotide flanked by any pair of nucleotides to change to any of the other three, is computed from all mutations observed in each cohort. For the benchmark described in this article, the mutational profile calculated within the intOGen pipeline (17) was used.

### 3D clustering score of individual residues

Given a protein-coding gene of interest alongside a matching protein conformation, a catalog of mutations mapping to that gene and a specific residue of interest, say *R*, we define a “clustering score” to reflect how extreme is the number of mutations mapping to either the residue or its neighbors in 3D space (within a sphere *S* centered at *R* with a predefined radius) with respect to the per-site background mutation probabilities that we can infer from the mutational profiles described in the previous section.

We model the number of mutations within the sphere *S* as being drawn from a binomial distribution whereby the probability to draw a single mutation from *S* is equal to the aggregate probability mass comprising all per-site probabilities of missense mutations involving the residues within *S*. Concretely, if the catalog of mutations mapping to the gene has *N* mutations, we model the number of mutations in *S* as binomially distributed with parameters *N* and *p*, where *p* is the sum of all the per-site probabilities of missense mutation sites involving the residues of *S*. Using the 96-channel mutational profile (see section “Mutational profile” above) along with DNA sequences of the transcript associated with the protein, we computed the per-site missense mutation probabilities by requiring that each site has a probability proportional to the corresponding 96-channel and that the sum of probabilities across all sites in the transcript is 1. In the calculation of the binomial parameter *p*, we used the contact maps (see section “Calculation of contact maps” above) to determine which missense mutation sites map to the residues within *S* and the per-site mutation probabilities.

We denote the probability mass of a given count *s* of mutations in *S* as *P(s)*, i.e.

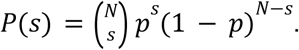

We can then compute 𝑃(𝑛 ≥ 𝑠) = 𝑃(𝑠) + 𝑃(𝑠 + 1) +…+ 𝑃(𝑁) the probability that *s* mutations or more had been drawn from *S* according to this binomial model. Intuitively, the smaller *P(s)* the more extreme the number of mutations mapping to *S*.

It is important to note that we aimed at a score that could be used to compute empirical p-values (as explained in the section below) for every gene, which we want to use to rank the genes by significance. While the log-probability -log 𝑃(𝑛 ≥ 𝑠) yields an intuitively sound measure, this score has the caveat that it does not cast a comparable distribution across genes with different *N* and as a result the significance that we can derive using this score ends up reflecting the order of magnitude of *N* (see section “Calculation of p-values” below). We thus defined the clustering score at residue *R* using the following likelihood ratio:

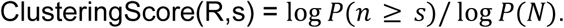

Finally, we normalized this 3D clustering score to enable comparisons across different genes and improve interpretability. Specifically, we divided the observed score by the average of the distribution of synthetic scores (plus one standard deviation unit) at the corresponding rank. The resulting score is proportional to the deviation from neutrality: a score of 1 indicates no deviation, while a score of 2 means the observed clustering signal is twice as high as expected under neutral mutagenesis. It is this normalized 3D clustering score (Supplementary Methods) that is shown in all Figures throughout the manuscript, while the non-normalized, previously described 3D clustering score is used to compute the p-values of mutated residues (see below).

### Calculation of p-values

Given a catalog of *N* observed missense mutations in a protein-coding gene, by using the method presented in the previous section we can endow each mutated residue with a clustering score, thereby giving a vector of scores that we can rank. To render an empirical p-value for each mutated residue, we build a distinct null model for each rank. For a residue *R* ranked n-th by clustering score, we compute an empirical p-value by comparing its (observed) clustering score with an empirical null distribution for the n-th rank. This empirical distribution is computed as follows: for *k=*10,000 iterations, we randomly draw *N* mutations with replacement among the available missense mutation sites in the protein-coding gene, with probability proportional to the relative mutation rate per trinucleotide context, given by the mutational profile. Each random catalog of mutations yields a vector of clustering scores associated with the mutated residues. For each such vector, we sort the entries in decreasing order, thus producing a “ranked vector” of clustering scores. By stacking the ranked vectors generated across *k*=10,000 iterations, we can build a null empirical distribution per rank by taking all the clustering scores that happen to occupy the query rank in the stack of ranked vectors.

For the residue in the *n*-th rank we compute an empirical p-value with a standard deviation offset, meaning that we categorize all null scores within one standard deviation of the observed score as equally extreme to the observed score. This approach is slightly more conservative than the common notion of empirical p-value definition and it is justified because of the propensity of the clustering score to form narrow groups, particularly when the number of mutations observed in the gene is low (see Supplementary Methods). We compute the empirical p-value associated with a residue *R* ranked n-th as *T(n), defined as* the proportion of values in the rank *n* null empirical distribution *D(n)* (plus one standard deviation unit) that are higher than the observed clustering score at *R* by more than one standard deviation of the null distribution,

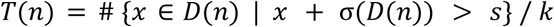

where σ*(D(n))* is the standard deviation of the empirical distribution *D(n)* and *s* is the observed clustering score at *R*. We set the p-value of a gene as the minimum empirical p-value reached across all mutated residues. Finally we correct for multiple testing (Benjamini–Hochberg FDR method) (44) across all the protein-coding genes analyzed to obtain a q-value per gene. By default, a residue is considered to be a significant mutational 3D cluster if its p-value is below 0.01, and a gene is considered significant if its q-value is below 1%.

### Agglomerating significant mutational 3D clusters into clumps

The fact that several residues are identified as significant by Oncodrive3D in the same protein may well be the manifestation of a common latent structural feature that is worth identifying; for example, functional domains or motifs that when mutated underlie the mechanism of tumorigenesis of the protein in question may be revealed by the significant 3D clusters mapping onto it. Thus, it is an important question whether the significant mutational 3D clusters found in a protein are structurally related. To answer this question, we grouped the significant 3D clusters into so-called “clumps” based on their structural closeness.

To do this, all residues identified as significant mutational 3D clusters by Oncodrive3D are used as nodes to build a graph. The edges of this graph represent the proportion of mutations “shared” between each pair of nodes, calculated as the ratio of the number of mutations within the residues shared by the volumes around the two residues and the total number of mutations associated with those residues. Nodes in the graph are then clustered using the NetworkX (45) implementation of the label propagation algorithm, a semi-supervised, network-based machine learning method for community detection (46).

### Oncodrive3D benchmark

To obtain metrics of sensitivity and specificity of driver discovery methods (including Oncodrive3D), we used two lists of *i) bona fide* cancer driver genes (positive instances) and *ii)* likely non driver genes which are often identified as false positives by driver discovery methods (see above). The overlap of a set of genes identified by a driver discovery method and the former list provides a measurement of its sensitivity, while its overlap with the latter constitutes a (reverse) measurement of its specificity.

We compared these measurements of sensitivity and specificity for Oncodrive3D across 215 cohorts of tumors (see above) and seven other state-of-the-art driver discovery methods: dNdScv (3), OncodriveFML (6), OncodriveCLUSTL (11), CBaSE (8), MutPanning (7), HotMAPS (10), SMregions (12). For each one of these methods, genes with p-value below 0.01 were considered significant. To rule out differences attributable to variation in the input data, we ran all methods in parallel on exactly the same list of mutations in every cohort. To do this, we took advantage of the fact that these seven state-of-the-art driver discovery methods are run by the intOGen pipeline (17), so that they receive exactly the same list of mutations as input. To run Oncodrive3D, we used this same list of mutations, that is after pre-processing by the pipeline (47) (e.g., filtering of hypermutated samples; see above). We also mapped mutations for Oncodrive3D to MANE Select transcripts for all AlphaFold models generated from these transcripts (17,334 proteins), or to ENSEMBL canonical transcripts for cases in which MANE Select transcripts were not available.

Still, comparing these measurements of sensitivity and specificity across methods is not a trivial task. First, by design, the methods do not process the exact same set of genes (e.g., some methods, such as HotMAPS and Oncodrive3D focus on genes affected by missense single nucleotide variants, while others also analyze genes affected by variants with other consequences). To avoid this hurdle, we compared the performance of methods on the set of genes processed by all of them. Secondly, different methods measure different signals of positive selection, using different statistical tests, and the list of significant genes according to various methods are thus not immediately comparable. To solve this problem, we reasoned that p-values produced by driver discovery methods can be used to rank the genes it identifies. We reasoned that –as we have done before (17)– a comparison of how their ranking of genes overlap with the positive and negative sets of genes, instead of a direct comparison of the list of significant genes produced by each, could solve this hurdle.

Briefly, to compare the sensitivity and specificity of several driver discovery methods, only genes processed by all methods are included, and only cohorts with at least 5 processed genes across all methods are analyzed. In the case of Oncodrive3D, only genes with more than two mutations –that is, in which a cluster could form– were processed. Ranked lists of these genes are then produced, based on the p-values awarded by each method. When p-values are identical, the effect size (i.e., the normalized 3D clustering score in the case of Oncodrive3D; see above) is used as a tiebreaker to determine the ranking. Each ranked list is examined one rank at a time, and the fraction of genes up to that rank that overlap with the positive (CGC) and negative (Fishy) lists of genes is annotated. Two curves are thus obtained, formed by these two fractions at every position of the ranking. These curves are cut at the rank *N*, defined as the minimum number of processed genes across all methods or 100, whichever is smaller for the cohort in question. The area under the curve is then calculated as

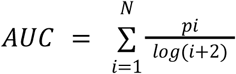

where *pi* is the enrichment score at any rank *i*.

For the sake of simplicity in the comparison, we make this AUC value relative to the maximum possible value (i.e., assuming that all genes up to rank *N* are included in the CGC or Fishy lists). This effectively re-scales the values of CGC-AUC and Fishy-AUC within the [0,1] interval. The re-scaled AUC is thus given by

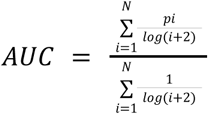

The area under the two resulting curves for each method are termed CGC-AUC (area under the curve formed by the fraction of genes within the CGC at each ranking position) and Fishy-AUC (area under the curve formed by the fraction of genes within the Fishy list at each ranking position). CGC-AUC constitutes a proxy measurement of the sensitivity of the method, while Fishy-AUC is a proxy measurement of its specificity. Therefore, we use CGC-AUC and Fishy-AUC to compare the sensitivity and specificity of seven state-of-the-art driver discovery methods and Oncodrive3D.

### Computational efficiency metrics

Taking advantage of the intOGen pipeline, we executed seven state-of-the-art driver discovery methods in parallel using Nextflow (48). Oncodrive3D was executed separately, also using Nextflow. Metrics for execution time and memory usage were extracted from the trace file (*trace.txt*) generated by Nextflow. We defined runtime as the actual execution time of each method (the *realtime* field in the trace), converted to hours, and multiplied by the number of CPUs utilized to calculate the total CPU-hours consumed by each method. Similarly, we defined memory usage as the peak virtual memory allocated to each method during execution (the *vmem* field in the trace).

### Structural models of significant 3D clusters

To visualize the recurrence of significant clusters across cohorts, protein structures were rendered using ChimeraX software (49). Each protein structure was displayed using a cartoon representation, which effectively highlights the secondary structure elements, such as alpha helices, beta sheets, and loops, and was uniformly colored in gray. Residues associated with significant clusters were annotated and highlighted using the atomic representation in spheres style, which displays the positions of individual atoms. These residues were colored according to the cluster recurrence across cohorts, where the color variation reflects the frequency at which the cluster was observed across the cohorts in which the corresponding gene was detected as significant.

### Annotation of genes identified by Oncodrive3D across cohorts of tumors and normal blood

Gene annotations presented in the heatmaps (Figures 5a, Supplementary Figures 9 and 12a), and used for the counts in Figure 5b, were obtained from the Cancer Gene Census (CGC) (42), OncoKB (43), and results obtained running the intOGen pipeline on the same cohorts. Specifically, genes were annotated as CGC if any of the tumor types listed in the “Tumor Types(Somatic)” column of the CGC file matched the tumor type of the corresponding cohort at the organ level, as shown in Supplementary Table 1. To map each tumor type to an organ or system of organs, we used the OncoTree (50). Specifically, for each cancer type, we traced its lineage up the tree until the point just before differentiation into solid and non-solid tumors.

### Code accessibility

Oncodrive3D can be freely downloaded, installed and run as a python standalone script (https://github.com/bbglab/oncodrive3d). A pre-built software container for Oncodrive3D can also be obtained from https://hub.docker.com/r/bbglab/oncodrive3d. Python notebooks ready to reproduce downstream analyses described in the paper are available at https://github.com/bbglab/oncodrive3d_paper.

## Results

### Oncodrive3D identifies mutational 3D clusters that deviate from neutrality

Oncodrive3D identifies significant deviations in the clustering of mutations in the 3D structure of a protein observed across tumors from that expected under neutrality in the same set of tumors. To that end, the local accumulation of observed mutations in the three-dimensional (3D) space (3D clustering score) at all mutant amino acid residues in the three-dimensional space is scored and compared to that computed for synthetic mutations generated following the process of neutral mutagenesis.

The input of Oncodrive3D is the list of non-synonymous mutations observed in each protein-coding gene across a cohort of tumors, as well as the trinucleotide profile of substitutions (obtained from all types of single nucleotide variants). This profile reflects the underlying mutational processes active throughout the life of the cells that form the tumor and can thus be used to model the neutral mutagenesis of the tissue in question. Oncodrive3D then maps the observed non-synonymous mutations to the 3D structure of every protein and counts all the mutations falling within spheres of a configurable diameter centered at each mutated amino acid residue (Fig. 1a). The configuration of the spheres takes into account the uncertainty of the location of residues within the AlphaFold model of the protein in question (Methods). Next, a 3D clustering score for each mutated residue is computed, on the basis of the probability (following a binomial distribution) of observing the same or more mutations within its corresponding sphere (Methods) given the total number of mutations observed across samples.

**Figure 1.**
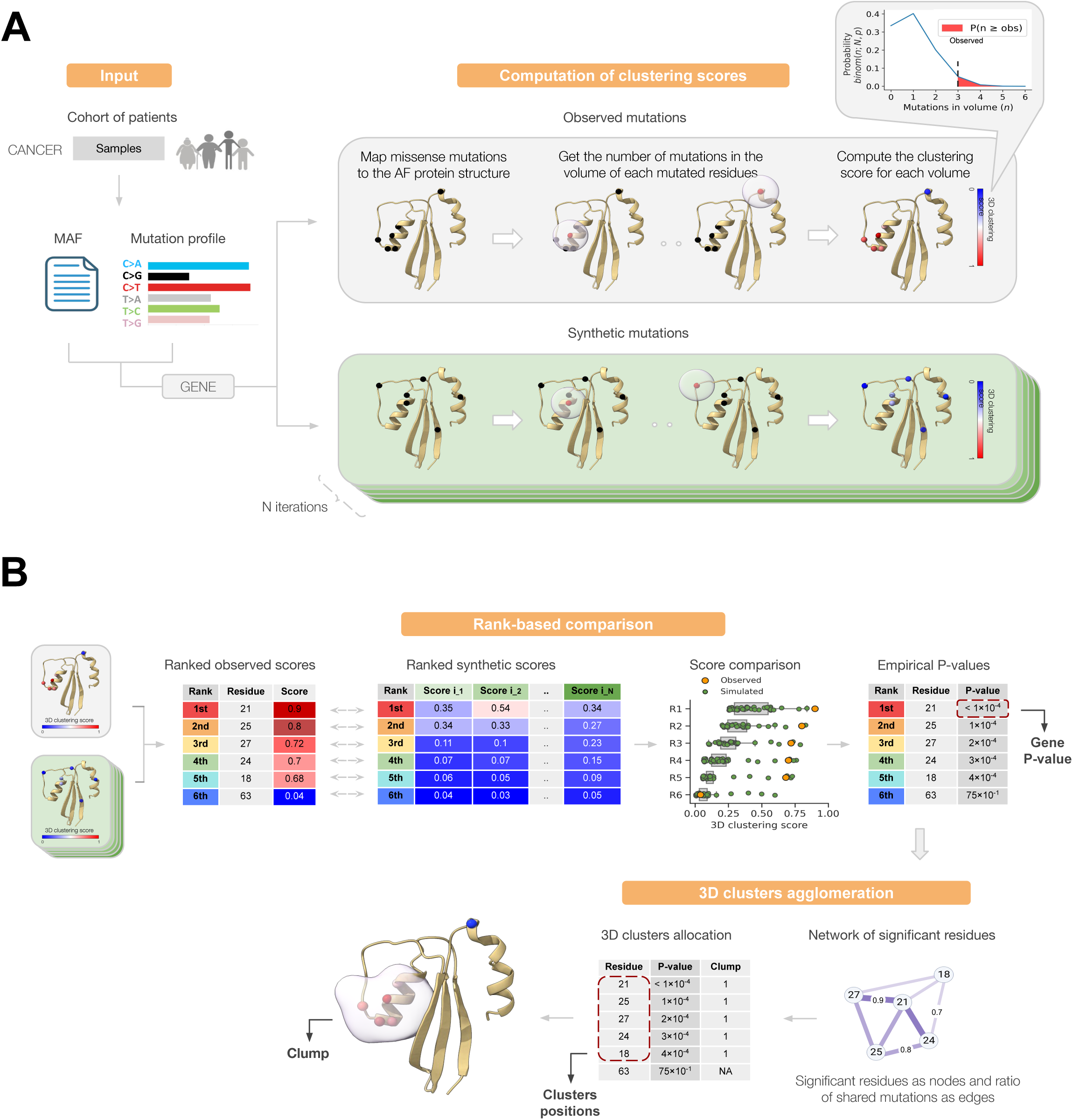
Oncodrive3D detects significant mutational 3D clusters in proteins. A) The left panel describes the input of Oncodrive3D (a list of missense somatic mutations in genes across a cohort of individuals), and the calculation of the profile of trinucleotide changes from this list of mutations. The right panel presents a schematic representation of the calculation of the observed (top) 3D clustering score for spherical volumes containing every mutated residue in a toy protein and of expected (bottom) 3D clustering score for spherical volumes containing residues with synthetic mutations. In 10,000 (or any predetermined number) iterations, synthetic mutations (the same number of observed mutations in every iteration) are generated following the probabilities of trinucleotide changes in the profile, as these represent the underlying process of neutral mutagenesis. B) The top panel describes the calculation of the significance of the observed 3D clustering score of every mutated residue, through a rank-based comparison with the expected 3D clustering scores computed on the basis of synthetic mutations. The bottom panel describes the agglomeration of all residues with significant mutational 3D clustering scores to obtain relevant clumps in the 3D structure of the protein. MAF = Mutation Annotation File, AF = AlphaFold

To compute the distribution of expected 3D clustering scores under neutral mutagenesis, the number of mutations observed in each gene are randomly distributed following the probabilities of the trinucleotide substitution profile, and this process is iteratively repeated (e.g. 10,000 times). In the example protein shown in Figure 1, at each iteration, synthetic 3D clustering scores for 6 residues are obtained. Carrying out 10,000 iterations yields a matrix of 60,000 (six rows by 10,000 columns) synthetic 3D clustering scores (Fig. 1b). Oncodrive3D then quantifies the significance of the deviation of the observed 3D clustering clustering scores of the six mutated residues from those calculated under neutrality. To this end, it compares the ranked observed 3D clustering scores of the six mutated residues to ranked synthetic 3D clustering scores, rather than a direct comparison of observed and simulated 3D clustering scores of each mutated residue. That is, the residue with the highest observed 3D clustering score is compared with the 10,000 top-ranking synthetic 3D clustering scores. This accommodates the possibility that amino acid residues with high clustering scores may be generated randomly through background mutagenesis, without any positive selection whatsoever. An empirical p-value is then computed, as the fraction of all elements of the distribution of synthetic 3D clustering scores equal to or greater than the observed clustering score of equivalent rank (Methods). Mutated residues with p-value below 0.01 are considered significant mutational 3D clusters in this analysis, and the p-value of the most significant residue is used to represent the gene after correction for multiple testing across all genes analyzed.

Next, Oncodrive3D agglomerates all significant mutational 3D clusters in a protein into clumps (higher-order clusters) of residues based on their proximity in space. To do this, all significant mutated residues are represented as nodes in a network, and each pair of nodes is connected if the spheres centered at the corresponding mutated residues share at least one mutation. The strength of the edge is proportional to the fraction of shared mutations between the two spheres. The nodes of the network are then agglomerated based on the strength of the connections between them (Methods). In the example illustrated in Figure 1b, the five significant mutational 3D clusters form a single clump in the protein. Thus, the output of Oncodrive3D consists of a list of proteins with a significant signal of 3D clustering of mutations across a cohort of tumors, as well as the location of clumps (agglomerated significant mutational 3D clusters) of each protein.

### Performance of Oncodrive3D across cohorts of tumors

Evaluating the specificity and sensitivity of driver gene discovery methods across a cohort of tumors of a given malignancy is not trivial, as no list of driver/non-driver genes in each tumor type is available. We and others have proposed that the manually curated genes in the Cancer Gene Census (CGC) (42) may be used as a general list (i.e. across different tumor types) of true driver genes. Comparing the output of a driver discovery method with this list may thus be used as a proxy measurement of sensitivity. Moreover, the comparison of the output with a list of genes with high mutation rate (due to their length, low expression, or other causes) across tissues (or Fishy genes; Supplementary Table 2), but with no evidence of involvement in tumorigenesis, may be used as a proxy of specificity of the method (see Methods).

The use of these two metrics is illustrated with a toy example of the results of a driver discovery method on a cohort of tumors (Fig. 2a, left). First, all genes bearing a significant p-value as per the method are ranked following their p-values. At each position in the ranking, the fraction of genes at the position or above annotated as *bona fide* cancer drivers in the CGC (orange points in Fig. 2a, right), or considered Fishy (green points), is computed. The resulting curves may be cut at any arbitrary position in the ranking and the areas below each of them (CGC-AUC and Fishy-AUC) up to that position, computed. In summary, the larger the computed CGC-AUC and the smaller the computed Fishy-AUC, the higher the sensitivity and specificity of the driver discovery method.

**Figure 2.**
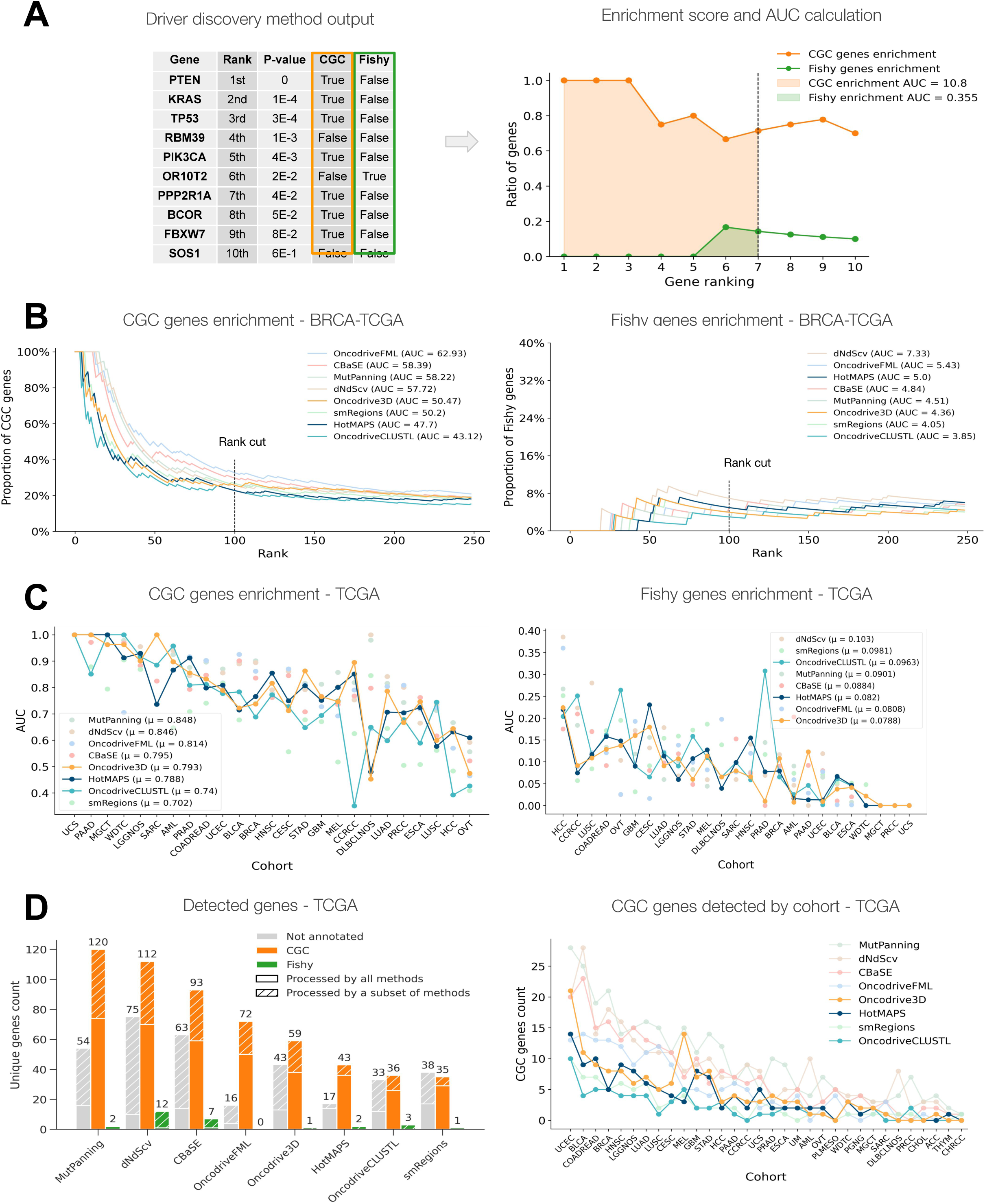
Performance of Oncodrive3D in comparison with other clustering-based driver discovery methods. A) Description of the estimation of the sensitivity and specificity of methods aimed at identifying signals of positive selection in the mutational patterns of genes using a toy example. The left panel presents a ranked list of genes (by their p-values) identified by a hypothetical method. Running down the ranking, at every position the fraction of genes of smaller or equal rank to the position in question that are either *bona fide* cancer genes (annotated in the Cancer Gene Census, or CGC) or likely false positives (in a manually curated list of “fishy” genes) is recorded and placed in a plot (right panel). The area under both resulting curves (AUC-CGC and AUC-Fishy) is highlighted in the corresponding color. B) CGC enrichment (sensitivity estimation) and Fishy enrichment (specificity estimation) curves for seven state-of-the-art driver discovery methods and Oncodrive3D on the TCGA-BRCA cohort. The corresponding AUC-CGC and AUC-Fishy are indicated. C) AUC-CGC and AUC-Fishy values of the same seven methods for 25 TCGA cohorts representing the same number of malignancies. The mean AUC-CGC and AUC-Fishy values for each method are indicated. D) The left panel presents the total number of CGC, Fishy and not-annotated (in either of these two lists) genes identified by seven state-of-the-art driver discovery methods and Oncodrive3D. The right panel presents the number of CGC genes identified by the seven methods in every TCGA cohort.

We are thus able to compare the performance of driver identification methods across any cohort of tumors, as exemplified through the TCGA breast adenocarcinomas cohort (BRCA, N=1,009). To compute the CGC-AUC and Fishy-AUC of the eight driver discovery methods illustrated in the figure (Fig. 2b), only genes analyzed by all methods are included, and the ranking for all methods is cut at position 100. In this cohort, Oncodrive3D shows greater CGC-AUC and smaller Fishy-AUC than HotMAPS and OncodriveCLUSTL, state-of-the-art driver discovery methods exploiting signals of mutational 3D clustering and linear clustering, respectively (Fig. 2b).

To systematically benchmark Oncodrive3D in comparison with other well-established driver discovery methods, we applied it to 32 TCGA cohorts, of the same number of malignancies (N=10,105 samples; Supplementary Table 1). The CGC-AUC and Fishy-AUC for the 25 cohorts with at least 5 genes analyzed by the eight benchmarked methods are shown in Figure 2c. We also employed a second larger dataset for this purpose, composed of 215 cohorts, including TCGA (intOGen cohorts; N=28,067 samples; Supplementary Fig. 1; Supplementary Table 1) (17). To facilitate the interpretation of CGC-AUC and Fishy-AUC, we calculated the highest possible values that can be obtained by applying each method to each cohort, and made the CGC-AUC and Fishy-AUC relative to those maximum values, effectively re-scaling them to values between 0 and 1. We used these relative CGC-AUC and Fishy-AUC values to compare the performance of Oncodrive3D to that of seven well-established mutational driver gene detection methods –dNdScv (3), OncodriveFML (6), OncodriveCLUSTL (11), CBaSE (8), MutPanning (7), HotMAPS (10), and smRegions (12)– which are implemented within the intOGen pipeline (17). Two of these methods –HotMAPS (10) and OncodriveCLUSTL (11)– exploit the clustering of mutations (in the 3D structure of the protein and its linear sequence, respectively), while the other five exploit other signals of positive selection, such as unexpected recurrence of protein-affecting mutations –dNdScv (3), and CBaSE (8)–, functional impact bias of mutations –OncodriveFML (6)–, enrichment of mutations in protein functional domains –SMRegions (12)–, and the deviation from neutral mutational trinucleotide contexts –MutPanning (7).

Across 25 TCGA cohorts (Supplementary Table 1) with at least five genes analyzed by all driver discovery methods used for benchmark (see Methods), Oncodrive3D outperforms HotMAPS and OncodriveCLUSTL in the CGC-AUC (0.793, 0.788 and 0.74; Fig. 2c, left). Oncodrive3D also outperforms HotMAPS and OncodriveCLUSTL on specificity, with the lowest Fishy-AUC (0.0788, 0.082 and 0.0963; Fig. 2c, right). Across the 215 cohorts in intOGen, similar results are obtained although, overall, the average CGC-AUC for HotMAPS is slightly higher than that of Oncodrive3D (Supplementary Fig. 2a).

In absolute numbers, Oncodrive3D identifies more CGC genes across 32 TCGA cohorts (with variable numbers of drivers identified per cohort) and across 215 intOGen cohorts than HotMAPS or OncodriveCLUSTL (Fig. 2d; Supplementary Fig. 2b). Moreover, it identifies more potentially novel driver genes across cohorts than HotMAPS and OncodriveCLUSTL. The increase with respect to HotMAPS may be explained, at least in part, by the fact that Oncodrive3D, by exploiting AlphaFold models, is able to analyze 4483 more genes –bearing at least two mutations in any cohort (see Methods)– across TCGA cohorts (5396 across 215 intOGen cohorts) than HotMAPS relying on experimentally solved 3D structures in the PDB. Nevertheless, when only genes analyzed by both methods are taken into account, Oncodrive3D still identifies more CGC genes and new potential drivers than HotMAPS. Some of these could be attributed to cases in which a cluster of mutations occurs in a region of the protein not covered by any experimental structure. We also verified that, in general, the distribution of p-values in the output of Oncodrive3D follows the expected uniform distribution, except for the few significant genes (Supplementary Fig. 3). This implies that the method is well calibrated for the tests carried out.

In summary, Oncodrive3D shows comparable performance to another state-of-the-art mutational 3D clustering method, and outperforms a linear mutational clustering method. The reliance of Oncodrive3D on AlphaFold models guarantees a maximum coverage of all potential cancer driver genes with significant mutational 3D clusters.

### Computational efficiency

Another important measure of performance of a computational tool is its efficiency, which can be measured in terms of CPU-hours and memory usage. This is particularly important for a tool intended for the systematic cumulative analysis of cohorts of tumors deposited in the public domain as part of the aim of constructing the compendium of mutational cancer driver genes (17). Therefore, we next compared the computational performance of Oncodrive3D and the other seven driver discovery methods implemented in the intOGen pipeline. We were particularly interested in comparing the computational efficiency of Oncodrive3D and HotMAPS, as they are both based on the same principle of identifying genes with significant mutational 3D clusters.

In terms of runtime, Oncodrive3D was among the fastest methods when run on TCGA cohorts (Fig. 3a). HotMAPS consumes 12 times more CPU-hours than Oncodrive3D to analyze the melanoma cohort. On the 32 TCGA cohorts, Oncodrive3D consumed 33 times less CPU-hours than HotMAPS (Fig. 3b). In terms of memory usage, Oncodrive3D also performed more efficiently than HotMAPS (1.19 TB vs 2.62 TB) in the analysis of the 28 TCGA cohorts (Fig. 3c,d). Similar differences were obtained in the analysis of all intOGen cohorts, with Oncodrive3D taking 38.6 times less CPU-hours than HotMAPS (Supplementary Fig. 4a). The memory usage of Oncodrive3D was also lower than that of HotMAPS (6.51 TB vs 16.6 TB) across intOGen cohorts (Supplementary Fig. 4b).

**Figure 3.**
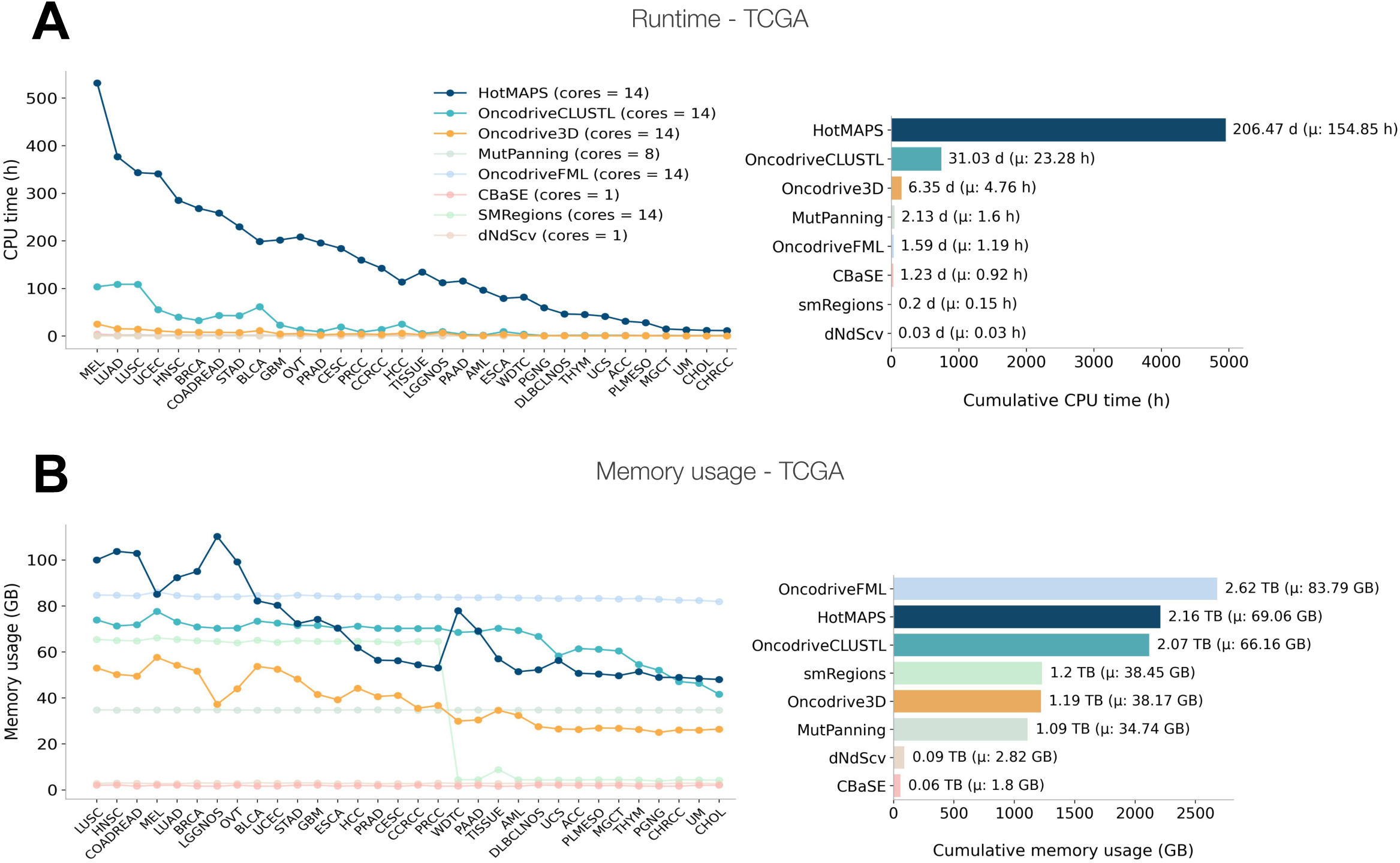
Computational efficiency of Oncodrive3D and other driver discovery methods. A) Efficiency of seven state-of-the-art driver discovery methods and Oncodrive3D in terms of CPU-time needed to process 32 TCGA cohorts. The left panel presents the number of CPU-hours used by each method to process each separate cohort, while the right panel presents the total number of CPU-days consumed by each method in the analysis of all cohorts. B) Efficiency of seven state-of-the-art driver discovery methods and Oncodrive3D in terms of usage of memory. The left panel presents the maximum GB used by each method to process each separate cohort, while the right panel presents the aggregated GB of memory used by each method in the analysis of all cohorts.

Thus, the computational efficiency of Oncodrive3D is superior to that of driver discovery methods based on mutational clustering. It performs on par with methods exploiting signals of positive selection that do not require operating with protein 3D structures.

### Complementarity with other driver discovery methods

We and others have previously shown that the identification of cancer driver genes is best approached through the combination of methods that exploit different signals of positive selection (14, 15, 17–19, 51, 52). Therefore, we next assessed the complementarity of Oncodrive3D and seven different driver discovery methods integrated within the intOGen pipeline.

Oncodrive3D identifies 103 genes across TCGA cohorts with significant mutational 3D clustering (Fig. 4a). Out of these, 30 are identified also by HotMAPS and OncodriveCLUSTL, with 39 more identified by one of them. This leaves 44 genes that are identified uniquely by Oncodrive3D as bearing a signal of mutation clustering, 16 of which are annotated in the CGC as *bona fide* cancer drivers (Fig. 4b). Only 20 out of these 44 are identified by some of the 5 remaining driver discovery methods implemented in intOGen, exploiting other signals of positive selection (Fig. 4c). Out of the remaining 24, which only Oncodrive3D identifies, five are annotated in the CGC (Fig. 4d). The proportion of genes uniquely identified by Oncodrive3D varies across TCGA cohorts (Fig. 4a-d).

**Figure 4.**
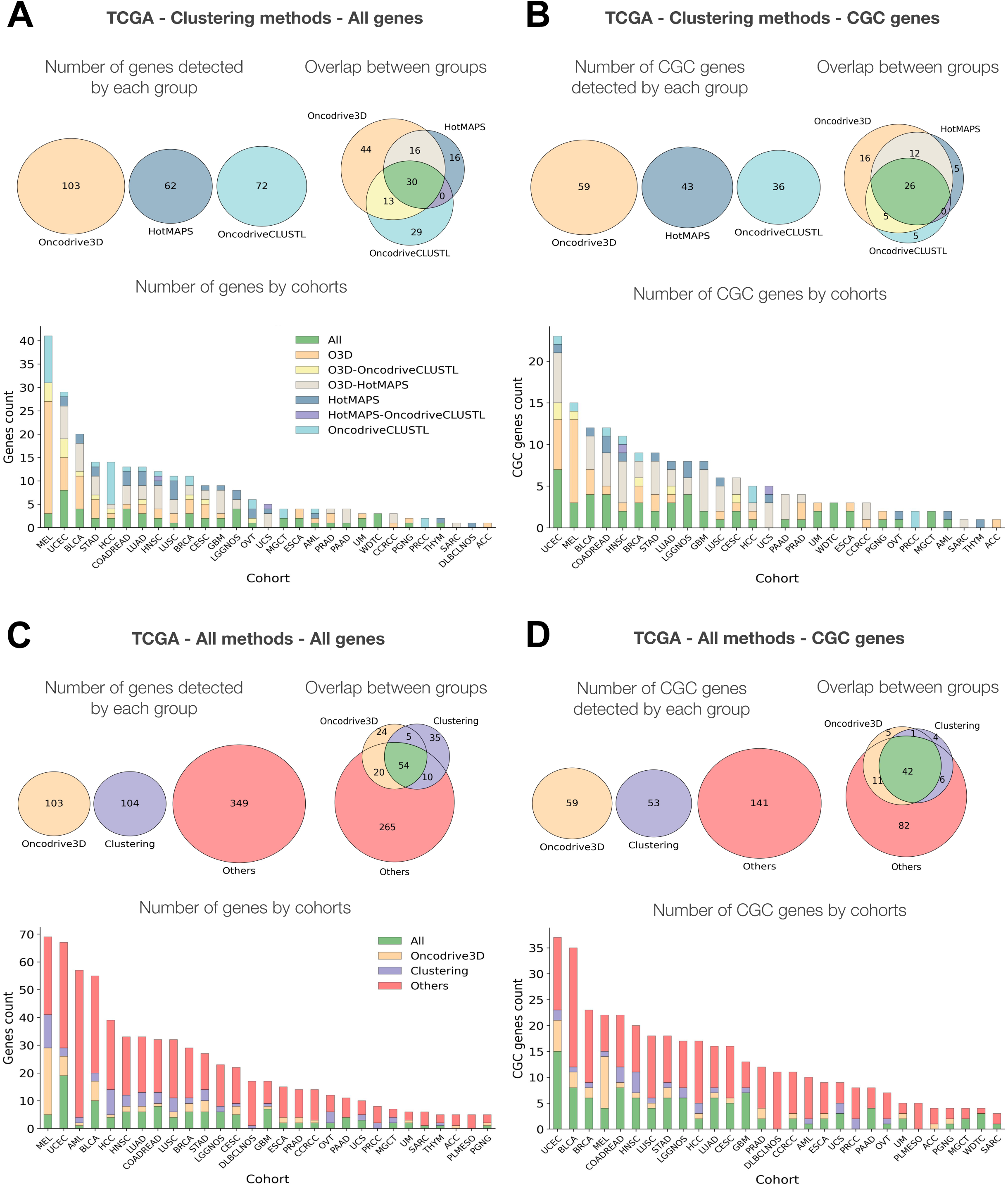
Oncodrive3D is complementary to seven other driver detection methods. A) Number of genes identified by two state-of-the-art driver discovery methods exploiting the clustering of mutations in the linear sequence of proteins (OncodriveCLUSTL) or in their 3D structure (HotMaps) and by Oncodrive3D. The overlap between genes identified by more than one method is represented for all TCGA cohorts through a Venn diagram (top panel) and for every cohort, through differently colored segments of bars. B) Same as A, restricted to *bona fide* cancer genes annotated in the CGC. C) Number of genes identified by seven state-of-the-art driver discovery methods and Oncodrive3D. State-of-the-art methods have been divided into two groups depending on whether they are based on detecting abnormal clustering of mutations in proteins (Clustering) or not (Others). The overlap between genes identified by methods in more than one group is represented for all TCGA cohorts through a Venn diagram (top panel) and for every cohort, through differently colored segments of bars. D) Same as C, restricted to *bona fide* cancer genes annotated in the CGC.

In all, 66 gene-tumor type combinations out of 190 identified by Oncodrive3D are missed by the two other clustering methods across TCGA cohorts (Supplementary Fig. 5a); 32 of these are missed when all seven methods currently included in intOGen are considered (Supplementary Fig. 5c). While 36 gene-cohort combinations involving genes annotated in the CGC were identified by Oncodrive3D, but not by the other two clustering methods (Supplementary Fig. 5b), 11 were missed by all methods in the intOGen pipeline (Supplementary Fig. 5d).

The same analysis across the 215 intOGen cohorts reveals similar results, with 132 (including 34 annotated in the CGC) out of 273 genes uniquely identified by Oncodrive3D that are missed by HotMAPS and OncodriveCLUSTL (Supplementary Fig. 6a,b), and 57 (4 in the CGC) missed by the seven methods in the intOGen pipeline (Supplementary Fig. 6c,d). Considering all gene-cohort combinations, 244 (including 130 with CGC genes) are missed by the two mutational clustering methods (Supplementary Fig. 6e,f), and 124 by all intOGen methods (31 with CGC genes) out of 619 identified by Oncodrive3D (Supplementary Fig. 6g,h).

These observations indicate that the results of Oncodrive3D complement well those of other driver discovery methods. Including it within the intOGen pipeline would thus contribute to the task of uncovering the compendium of mutational cancer driver genes across tumor types.

### Cancer driver genes with mutational 3D clustering signal

We next applied Oncodrive3D to somatic mutations identified across 32 TCGA cohorts and 215 intOGen cohorts of tumors. As expected, CGC genes present significantly higher 3D clustering scores than Fishy genes, with well-known oncogenes and TP53 exhibiting some of the highest values observed across all genes (Supplementary Fig. 7a). Interestingly, the distribution of scores of potential novel driver genes is similar to that of *bona fide* cancer driver genes (Supplementary Fig. 7b). Conversely, the 3D clustering scores of those deemed not significant closely resemble the distribution obtained for Fishy genes (Supplementary Fig. 7b). While clumps (aggregated significant mutational 3D clusters) in tumor suppressor genes predominantly occur in buried regions of the protein structure, those in oncogenes exhibit a bimodal distribution, with clumps found in both buried and exposed regions (Supplementary Fig. 7c, top left panel). Most clumps are located in rigid portions of the protein backbone (Supplementary Fig. 7c, top right panel) and generally involve regions of the protein structure with higher confidence scores from the AlphaFold model (Supplementary Fig. 7d, top right panel). As expected, clumps in tumor suppressors contain mutations that are more structurally destabilizing compared to those in oncogenes (Supplementary Fig. 7d, bottom left panel). Finally, clumps in oncogenes tend to harbor a higher number of per-residue mutations compared to those in tumor suppressors (Supplementary Fig. 7d, bottom right panel). Interestingly, potential novel driver genes (genes not annotated in the CGC or Fishy lists) show clump characteristics—such as solvent accessibility, AlphaFold local confidence (pLDDT; which in many cases serves as a proxy for backbone rigidity (53)), and stability changes—that align more closely with those of oncogenes than tumor suppressors, suggesting the prevalence of a gain-of-function-like mode of action among these potential new cancer drivers.

Out of 103 genes identified by Oncodrive3D as bearing significant mutational 3D clusters across 32 TCGA cohorts (Fig. 2d; Fig. 4a), 26 appeared significant in two or more cohorts (Fig. 5a; Supplementary Table 3). Most of these 26 genes are annotated in both, the CGC and OncoKB and they were identified by most or all driver discovery methods included in the intOGen pipeline. The number of genes with significant mutational 3D clusters was between 31 (melanoma) and 1 (adrenocortical carcinomas). These appear distributed differently among those annotated in the CGC for the tumor type in question, in other tumor types, or not annotated in the CGC, across cohorts. In all, out of 190 gene-TCGA cohorts identified by Oncodrive3D, 66 involve genes annotated in the CGC for the tumor type in question, 78 are associated to genes annotated in the CGC for other tumor types, while 46 correspond to novel candidate driver genes (Fig. 5b).

**Figure 5.**
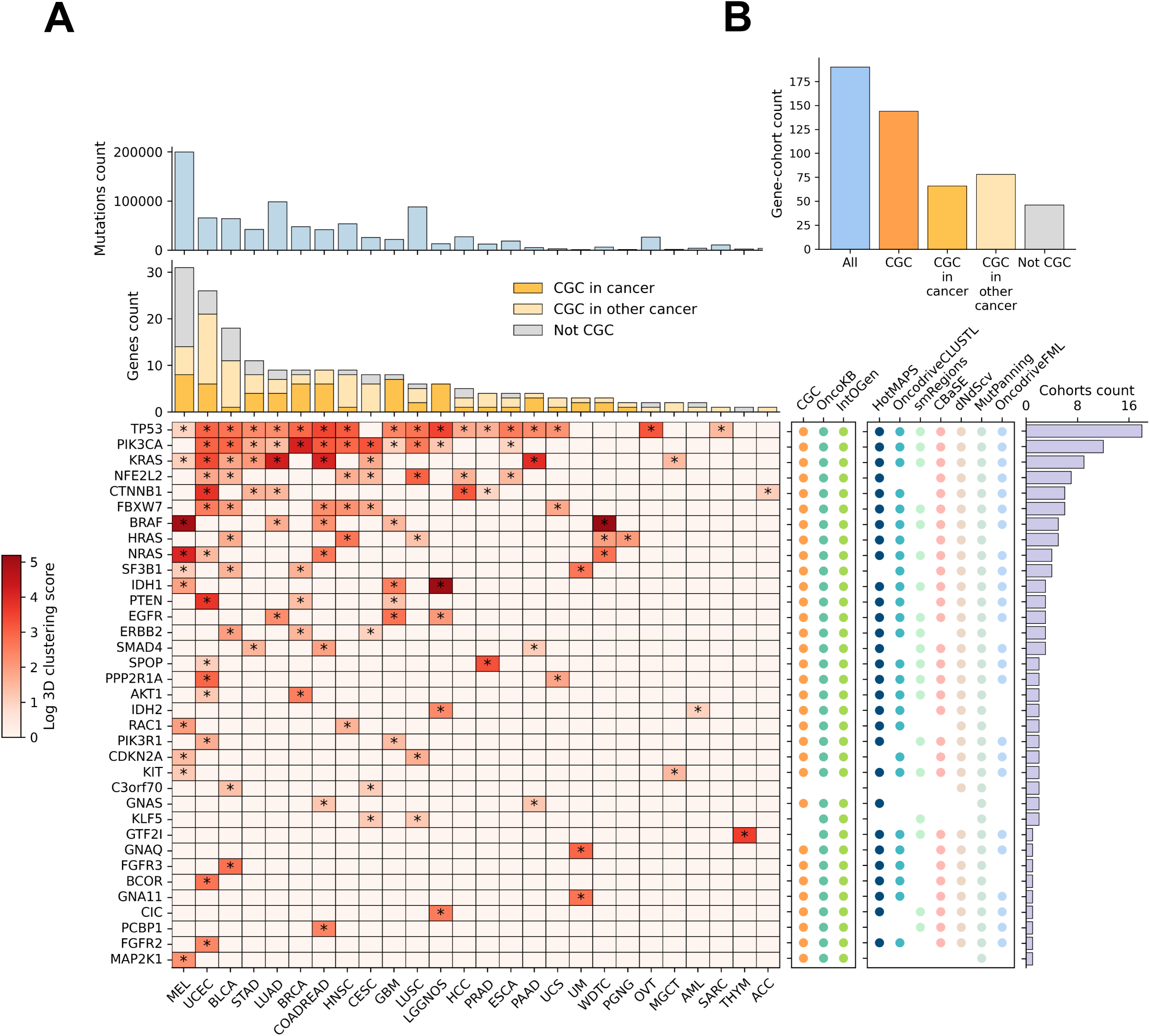
Oncodrive3D significant genes across TCGA cohorts. A) The central heatmap presents the top 35 significant genes (in terms of number of TCGA cohorts where they are identified by Oncodrive3D) according to Oncodrive3D (cells with an asterisk). The color of the cells represent the 3D clustering score of the residue with the lowest p-value in each gene. The two bar plots above the heatmap represent the total number of missense mutations in each cohort and the total number of genes (annotated in the CGC for the tumor type of the cohort in question, for other tumor types, or not annotated in the CGC) identified as bearing significant mutational 3D clusters by Oncodrive3D. The first rectangular panel by the right side of the heatmap denote whether the gene is annotated in any of two catalogues of *bona fide* cancer genes (CGC, OncoKB), or has been identified by the intOGen pipeline (17). The second rectangular panel denotes which of the driver discovery methods in the intOGen pipeline has identified the gene as a potential cancer driver. The bars at the right represent the number of TCGA cohorts where each gene has been found to bear significant mutational 3D clusters by Oncodrive3D. In the heatmap only 26 TCGA cohorts (those for which at least one driver gene is identified by Oncodrive3D) are included. B) Total number of gene-cohort combinations identified as bearing significant mutational 3D clusters by Oncodrive3D.

Across 215 intOGen cohorts, grouped by the organ or system of organs affected by the malignancy, a comparable landscape emerges (Supplementary Fig. 8a,b). The list of genes at the top of the heatmap (those with significant clusters across more organs) are the same as those at the top in the analysis of TCGA cohorts. Out of 619 gene-intOGen cohorts identified by Oncodrive3D, 204 correspond to CGC genes annotated to tumor types associated to the organ in question (see Methods), 215 involve genes annotated in the CGC for other tumor types, while 190 are potentially novel candidate driver genes (Fig. 5b). Looking at the other tail of the distribution, that is, genes with significant mutational 3D clusters in malignancies affecting only one cohort, we still found well-known CGC annotated drivers (Supplementary Fig. 9).

### Mutational 3D clusters under positive selection in cancer genes across tumor types

A second important output of Oncodrive3D is the location of significant mutational clusters in the tree-dimensional structure of cancer proteins. This provides information about the perturbation that the biological function of these genes suffer upon tumorigenesis in different tissues. This is illustrated through examples in Figure 6a,b and Supplementary Figure 10a-c. In NFE2L2, two clumps of significant mutational 3D clusters detected across 10 intOGen cohorts, representing tumors affecting 7 organs, (the first spanning across amino acid residues 24, 26-32, and 34, and the second across residues 77, and 79-82), overlap two degrons recognized by KEAP1, a joint adaptor and substrate receptor of a cognate E3-ligase (12, 54) (Fig. 6a). The recurrence of residues with mutational 3D clusters is variable. EGFR provides a very different example, with significant mutational 3D clusters detected at different locations of the protein in cohorts from different organs (Fig. 6b). These probably represent different mechanisms of tumorigenesis in these tissues.

**Figure 6.**
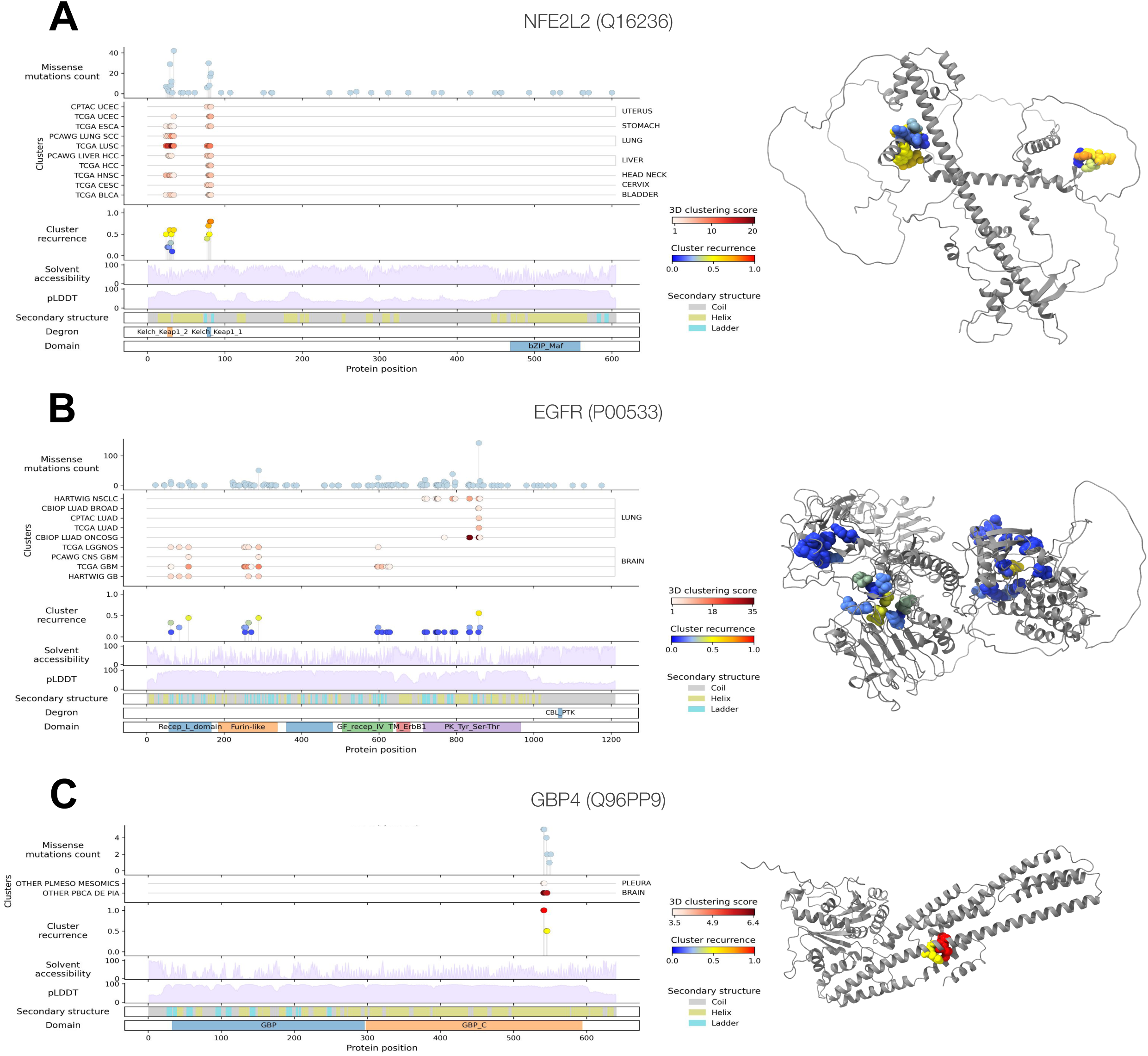
Recurrence of residues with significant mutational 3D clusters across cohorts of tumors. A) Recurrence of residues with significant mutational 3D clusters in NFE2L2 across 10 cohorts of tumors (annotated at the left of the second track) of 7 different organs (annotated at the right of the second panel). B) Recurrence of residues with significant mutational 3D clusters in EGFR across 9 cohorts of tumors of 2 different organs. C) Recurrence of residues with significant mutational 3D clusters in GBP4 across 2 cohorts of tumors of 2 different organs. In the three graphs, the top track presents the number of mutations affecting each residue of the protein. The second track presents the residues identified as bearing significant mutational 3D clusters by Oncodrive3D in each cohort of tumors analyzed, with the color representing the 3D clustering score of each of them. The third track represents the recurrence of each cluster (fraction of cohorts analyzed where the cluster is significant). The 4 tracks below present annotations of the protein structure that support the interpretation of the functional relevance of the clusters; from top to bottom: solvent accessibility, AlphaFold local model confidence (proxy for backbone rigidity), secondary structure and functional domains.

We also explored examples of genes with significant mutational 3D clusters which can represent novel driver genes. GBP4, for instance, encodes a guanylate-binding protein, a family known to be capable of promoting inflammation (55). Significant mutational 3D clusters identified in this protein appear partially conserved across tumors of the brain and pleura, and overlap its proposed C-terminal GTPase effector domain (56) (Fig. 6c). H3-3A is also identified as a potential novel driver gene with significant mutational 3D clusters in brain tumors (Supplementary Fig. 11b). It encodes a variant of one of the core histones (H3.3), some of which are *bona fide* cancer drivers via mutations affecting their methylation and, ultimately, the correct regulation of gene expression (57). Finally, Supplementary Figure 11c shows the recurrence of residues with significant 3D clusters identified in ESX1 across liver and pancreatic tumors. This gene encodes for a homodeomain transcription factor which has been proposed to act as a transcriptional repressor of KRAS (58).

### Application of Oncodrive3D to the identification of drivers of clonal hematopoiesis

We applied Oncodrive3D to the identification of genes with significant mutational 3D clusters in clonal hematopoiesis (CH), that is the normal expansion of clones in the blood. Analyzing the somatic mutations identified in the blood of 36,461 donors across three cohorts (29), Oncodrive3D identified several well-established CH driver genes (Supplementary Fig. 12a). Complementarity between methods is also important in the identification of CH driver genes (Supplementary Fig. 12b). A comparison of the mutational 3D clusters identified in DNMT3A in CH and those across AML cases revealed that significant residues within the methylase domain are more recurrent than those in other areas of the protein (Supplementary Fig. 12c). In the case of SF3B1, significant clusters across both CH and AML are confined to a small portion of the protein (Supplementary Fig. 12d).

## Discussion

Here, we have introduced Oncodrive3D, a novel computational method that identifies genes with significant mutational 3D clusters across tumors. Exploiting this signal of positive selection in tumorigenesis, Oncodrive3D detects genes capable of driving tumorigenesis. Moreover, the location of clumps of significant mutational 3D clusters detected in genes provide relevant information about their mechanisms of tumorigenesis in different tissues.

The annotation of significant mutational 3D clusters will also contribute to the identification of driver mutations across cancer genes which is carried out by machine learning-based models (25). With the use of AlphaFold 2 models (28), the annotation of mutational 3D clusters will cover all mutations in driver genes, thus extending the discovery of driver mutations.

Oncodrive3D shows sensitivity and specificity on par with another state-of-the-art method of detection of mutational 3D clusters (10), and outperforms a method aimed at detecting linear clusters (11). It clearly outperforms these two clustering-based driver discovery methods in the efficiency of the use of computational resources. Moreover, the results of Oncodrive3D are complementary to those obtained by other driver discovery methods based on other signals of positive selection (3, 6–8, 10–12). We thus plan to include it within the intOGen pipeline (17), which combines the output of seven driver discovery methods and is systematically run on cohorts of tumors collected from the public domain to produce the compendium of mutational driver genes. Oncodrive3D is available for local installation to run as a highly-configurable standalone method at https://github.com/bbglab/oncodrive3d. Oncodrive3D could be extended in the future to include the identification of mutational 3D clusters in the interface between interacting proteins. In summary, we envision that Oncodrive3D will make a very important contribution to the discovery of cancer driver genes and mutations, and the elucidation of the mechanisms of tumorigenesis of cancer genes across tissues.

## Data Availability

All cancer genomics datasets used in this study to test Oncodrive3D are deposited in the public domain. For example, somatic mutations identified in tumors of 32 cohorts by TCGA can be accessed via the Genomic Data Commons (GDC) (59). A list with the 215 cohorts employed here and their provenance can be found at Supplementary Table 1 and https://www.intogen.org.

## Acknowledgements

The authors wish to thank Federica Brando for her contribution to reviewing, organizing and packaging the code; and Ferriol Calvet for his contribution to the development of the method. The authors wish to acknowledge the donors and their families for generously contributing their tumor samples. Institute for Research in Biomedicine Barcelona is a recipient of a Severo Ochoa Centre of Excellence Award from the Spanish Ministry of Economy and Competitiveness (MINECO; Government of Spain) and an Excellence Institutional grant by the Asociación Española contra el Cáncer, and is supported by CERCA (Generalitat de Catalunya).

