## Supplemental Material for "Oncodrive3D: Fast and accurate detection of structural clusters of somatic mutations under positive selection"

### Table of Contents

**Supplementary Methods** **3**
Calculation of normalized 3D clustering scores and p-values 3
Oncodrive3D input and output 5
**Supplementary Note** **11**
Characterization of mutational 3D clusters 11
**Supplementary Figures** **13**
**Supplementary Tables description** **39**
**References** **40**

### Supplementary Methods

#### Calculation of normalized 3D clustering scores and p-values

To assess the significance of mutational 3D clusters, we compared their observed 3D clustering score with the distribution of synthetic 3D clustering scores of equivalent rank (see main manuscript Methods and Results and Figure 1 of these Supplementary Methods). We thus computed a so-called *normalized 3D clustering score* that puts the observed 3D clustering score of the residue within the context of its expectation. To that end, we divided the observed 3D clustering score by the mean of the distribution of synthetic 3D clustering scores plus one standard deviation (Figure 1). The resulting normalized 3D clustering score represents the factor whereby the observed score exceeds the upper standard deviation boundary of the distribution of synthetic 3D clustering scores. This normalization enables more consistent comparisons of clustering scores across residues within a gene and between different genes.

Resorting to the same rationale, we calculated an empirical p-value for each mutational 3D cluster within a gene (see main manuscript Methods and Results). To compute this p-value, we first compare the observed 3D clustering score of the residue to the distribution of synthetic 3D clustering scores of the corresponding rank after adding one standard deviation to all synthetic values. The p-value is then computed as the fraction of synthetic 3D clustering scores that exceed the observed 3D clustering score. This calculation makes the method more stringent than the conventional empirical p-value, ensuring that only clustering scores consistently higher than the typical synthetic 3D clustering score are considered noteworthy (Figure 1).

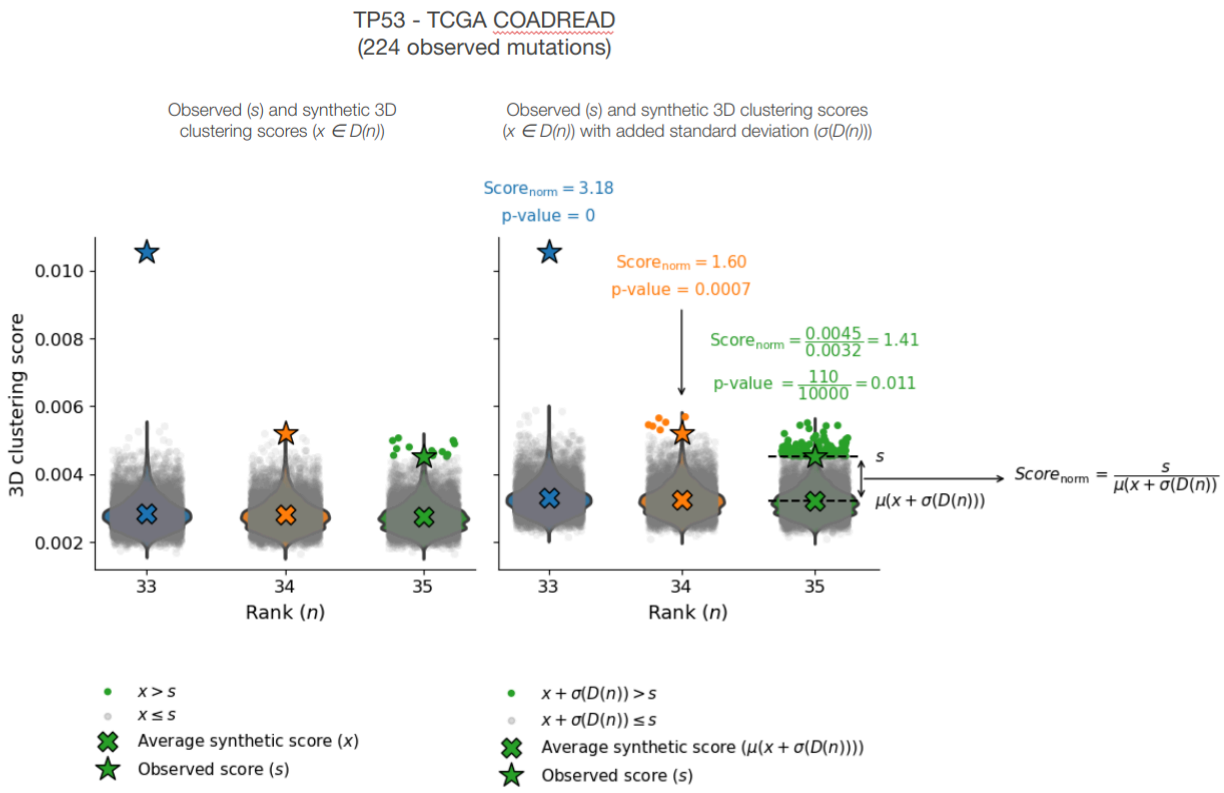


**Figure 1. Figure explaining the calculation of normalized 3D clustering score of mutational 3D clusters in a protein.**

The three residues occupying ranks 33, 34 and 35 of TP53 in the TCGA cohort of colorectal tumors are shown, with the stars at the values of their respective observed 3D clustering scores. The dots represent individual synthetic 3D clustering scores generated by separate iterations, with their distribution depicted by the corresponding scatter plots, and their mean as Xs. Synthetic 3D clustering score values equal to or greater than the observed value appear in blue, orange or green color (depending on the rank of the residue), while the remaining appear colored gray. In the left plot, the original synthetic scores, directly generated across iterations are shown.

In the right plot, the standard deviation of the distribution of synthetic 3D clustering scores at each rank is added to each point of the distribution, which is consequently shifted upwards. The normalized 3D clustering score of each residue results from dividing its observed 3D clustering score by the mean of this shifted distribution. The p-value of the cluster detected at each residue is calculated as the fraction of synthetic 3D clustering scores (after the shift resulting from adding the standard deviation of the distribution) that are equal to or greater than the observed 3D clustering score for the cluster in question.

#### Oncodrive3D input and output

The input of Oncodrive3D is the list of mutations observed in a gene across a cohort of samples. These mutations must be annotated with their consequence types, as Oncodrive3D considers only missense mutations for the 3D clustering analysis. Also, missense mutations need to be mapped to the protein coordinates of the MANE Select—or canonical (see main manuscript Methods)—transcripts of the genes. Both the MAF format (https://docs.gdc.cancer.gov/Data/File_Formats/MAF_Format/) or the output of the Ensembl VEP (1) can thus be used as input. Figure 2, below presents an example of an Oncodrive3D mutation input file in MAF format. Oncodrive3D also receives the mutational profile observed across all samples in the cohort. This is computed from the genomic immediate context of mutations of all consequences using a tool like SigProfilerMatrixGenerator (2) or bgsignature (https://pypi.org/project/bgsignature/) (Table 1). The output of the Ensembl VEP (1) can be also used as input for Oncodrive3D. This option has the advantage of containing the mapping of mutations to all transcripts of a gene, thus maximizing the number of matching transcripts between the input and the AlphaFold database (see Methods).

**Table 1. Example of Oncodrive3D input files in MAF format.**

| Hugo_Symbol | Reference_Allele | Tumor_Seq_Allele2 | Variant_Classification | Transcript_ID | HGVSp_Short |
|---|---|---|---|---|---|
| OR4F5 | G | T | missense_variant | ENST00000335137 | p.G145C |
| KLHL17 | G | A | missense_variant | ENST00000338591 | p.V133M |
| KLHL17 | G | C | missense_variant | ENST00000338591 | p.D402H |
| KLHL17 | G | A | synonymous_variant | ENST00000338591 | . |
| PLEKHN1 | C | T | stop_gained | ENST00000379410 | . |
| PLEKHN1 | G | T | missense_variant | ENST00000379410 | p.G152W |

The main output of Oncodrive3D consists of two files: a gene-level output (with extension `3d_clustering_genes.csv`) and a residue-level output (with extension `3d_clustering_pos.csv`), both summarizing the results of the 3D clustering analysis. The gene-level file provides a list of genes with mutations from the input file, including those not analyzed due to having only one or two mutations. It includes the 3D clustering score and p-value for the highest-scoring mutational 3D cluster (used as the gene's p-value), the corrected p-value, the coordinates of all significant 3D mutational clusters (and their aggregation into clumps) identified in the gene, a range of annotations for these clusters, and additional information. The residue-level file contains information for each mutational cluster, such as specific residues involved in clustering, the number of mutations within the volume of each mutated residue, and other related metrics. (Tables 2 and 3, below). Comprehensive details on the input and output formats are provided in the documentation (https://github.com/bbglab/oncodrive3d).

**Table 2. Example of Oncodrive3D gene-level output file.**

| Gene | Uniprot_ID | pval | qval | C_gene | C_pos | C_label | Score_ obs_sim_top_vol | Clust_res | Mut_in_gene | Clust_mut | Mut_in_top_vol | Mut_in_ top_cl_vol |
|---|---|---|---|---|---|---|---|---|---|---|---|---|
| PIK3CA | P42336 | 0 | 0 | 1 | [1049 1047 ..] | [0 0 ..] | 60.34 | 13 | 360 | 302 | 151 | 151 |
| TP53 | K7PPA8 | 0 | 0 | 1 | [238 194 ..] | [0 0 ..] | 17.94 | 65 | 215 | 211 | 72 | 149 |
| CTCF | P49711 | 0.0003 | 0.42 | 0 | [283 288 284] | [0 0 0] | 2.435 | 3 | 12 | 5 | 5 | 5 |

Note that the third gene in the exemplary table (CTCF), although bearing a significant mutational 3D cluster (p-value=0.0003), in the cohort shown in the table does not appear significant after the correction for false discovery (q-value=0.42).

Description of table columns:

- **Gene**: HUGO symbol or identifier of the gene being analyzed.
- **Uniprot_ID**: Identifier for the gene's protein product in the UniProt database.
- **pval**: The p-value indicating the statistical significance of the gene in the 3D clustering analysis (lowest p-value among the residues of the gene).
- **qval**: Adjusted p-value (q-value) to control for false discovery rate (FDR) using the Benjamini-Hochberg method.
- **C_gene**: Binary label indicating if the gene is detected to be significant (by default if q-value < 0.01).
- **C_pos**: List of protein positions of the gene clusters.
- **C_label**: List of labels indicating the clump to which each cluster is grouped.
- **Score_obs_sim_top_vol**: Normalized 3D clustering score of the gene (normalized 3D clustering score of the residue with the lowest p-value; if multiple residues have the same lowest p-value, the maximum normalized 3D clustering score among those residues is used).
- **Clust_res**: Number of residues detected as significant clusters.
- **Mut_in_gene**: Number of missense mutations in the gene.
- **Clust_mut**: Number of missense mutations in significant clusters.
- **Mut_in_top_vol**: Number of missense mutations in the most significant cluster (in the volume of the residue with the lowest p-value).
- **Mut_in_top_cl_vol**: Number of missense mutations in the clusters of the most significant clump (in the volume of residues of the clump that includes the residue with the lowest p-value).

Only a subset of the columns from the output is included here. For a more comprehensive and detailed description of the output, please refer to the Oncodrive3D documentation (https://github.com/bbglab/oncodrive3d).

**Table 3. Example of Oncodrive3D residue-level output file.**

| Gene | Uniprot_ID | Pos | Mut_in_gene | Mut_in_res | Mut_in_vol | Score | Rank | Score_obs_sim | pval | C | C_ext | Clump | Mut_in_cl_vol | PAE_vol |
|---|---|---|---|---|---|---|---|---|---|---|---|---|---|---|
| CTCF | P49711 | 283 | 12 | 1 | 5 | 0.271 | 0 | 2.267 | 0.0008 | 1 | 0 | 0 | 5 | 1.2 |
| CTCF | P49711 | 288 | 12 | 1 | 5 | 0.27 | 1 | 2.387 | 0.0005 | 1 | 0 | 0 | 5 | 1.8 |
| CTCF | P49711 | 284 | 12 | 3 | 5 | 0.262 | 2 | 2.435 | 0.0003 | 1 | 0 | 0 | 5 | 0.6 |
| CTCF | P49711 | 228 | 12 | 1 | 2 | 0.091 | 3 | 0.9 | 0.715 | 0 | | | | |
| CTCF | P49711 | 226 | 12 | 1 | 2 | 0.09 | 4 | 0.929 | 0.6628 | 0 | | | | |
| CTCF | P49711 | 443 | 12 | 2 | 2 | 0.088 | 5 | 0.958 | 0.6073 | 0 | | | | |
| CTCF | P49711 | 258 | 12 | 1 | 1 | 0.047 | 6 | 0.526 | 1 | 0 | | | | |
| CTCF | P49711 | 687 | 12 | 1 | 1 | 0.044 | 7 | 0.519 | 1 | 0 | | | | |
| CTCF | P49711 | 378 | 12 | 1 | 1 | 0.031 | 8 | 0.38 | 1 | 0 | | | | |

Note that the rank (*Rank)* is defined using the 3D clustering score (*Score*) assigned to each mutated residue, and the p-value (*pval*) and normalized 3D clustering score (*Score_obs_sim*) are computed at the rank level. As shown in the example, a lower rank does not always correspond to a lower p-value or a higher normalized 3D clustering score.

Description of table columns:

- **Gene**: HUGO symbol or identifier of the gene being analyzed.
- **Uniprot_ID**: Identifier for the gene's protein product in the UniProt database.
- **Pos**: Coordinate of the mutated protein residue.
- **Mut_in_gene**: Number of missense mutations in the gene.
- **Mut_in_res**: Number of missense mutations in the residue.
- **Mut_in_vol**: Number of missense mutations in the volume of the residue.
- **Score**: 3D clustering score for the residue.
- **Score_obs_sim**: Normalized 3D clustering score for the residue.
- **pval**: The p-value of the residue in the 3D clustering analysis.
- **C:** Binary label indicating whether the cluster at that residue is significant (1) or not (0). A cluster is marked as significant either because it meets the significance criteria directly or because it has been rescued by contributing mutations to another significant cluster.

- **C_ext:** Binary label indicating whether the cluster has been rescued by contributing mutations to another significant cluster (1) or if it was significant on its own (0).
- **Clump**: Identifier for the clump to which the cluster at that residue has been assigned.
- **Rank**: Rank used to perform the calculation of the normalized 3D clustering score and p-values.
- **Mut_in_cl_vol**: Number of missense mutations in the clusters of the clump to which the cluster at that residue has been assigned.
- **PAE_vol**: Weighted average predicted aligned error (PAE) of the residues in the volume of the cluster.

Only a subset of the columns from the output is included here. For a more comprehensive and detailed description of the output, please refer to the Oncodrive3D documentation (https://github.com/bbglab/oncodrive3d).

### Supplementary Note

#### Characterization of mutational 3D clusters

We explored the properties of 3D mutational clusters, focusing on their recurrence across tumors, their 3D clustering score distributions, and their structural characteristics across different gene annotations and modes of action.

We first analyzed the recurrence of significant mutational 3D clusters found in well-established cancer genes (Fig. 6 and Supplementary Fig. 10), potential novel driver genes (Fig. 6 and Supplementary Fig. 11), and well-established clonal hematopoiesis driver genes (Supplementary Fig. 12). In some genes, we observed that the same residues are consistently present across significant mutational 3D clusters. This is, for example, the striking case of TP53, where missense mutations in significant 3D clusters clearly concentrate in the P53 DNA binding domain of the protein (Supplementary Fig. 10a). The extension of the large clumps in which these TP53 mutational 3D clusters accumulate contrasts with very narrow clumps detected in several oncogenes across cancer types. For example, in the case of NFE2L2 (Fig. 6a), two handfuls of residues form two distinct clumps of mutational 3D clusters that perfectly overlap with two degrons (degradation signals) of this gene, resulting in the abnormal stabilization of the protein. These are very well conserved across all cancer types where NFE2L2 is found to be a driver. EGFR, on the other hand, presents a case in which clumps of significant mutational 3D clusters are organ-dependent, overlapping distinct domains of the protein in tumors originating in different parts of the brain or in the lungs.

We also studied the distribution of 3D clustering scores, defined for each gene as the 3D clustering score of its most significant residue, across CGC genes, Fishy genes, and non-annotated genes, distinguishing between those detected as significant by Oncodrive3D and non-significant ones (Supplementary Fig. 7a,b). The distribution of 3D clustering scores in *bona fide* cancer driver genes is significantly shifted towards high values (p-value=$2.5 \times 10^{-20}$) when compared to that of 3D clustering scores in Fishy genes (Supplementary Fig. 7a). The distribution of 3D clustering scores in other significant genes overlaps with that of *bona fide* cancer genes, whereas the distribution of 3D clustering scores in non-significant genes aligns with that of Fishy genes (Supplementary Fig. 7b).

Moreover, we analyzed the characteristics of clumps identified by Oncodrive3D across oncogenes, tumor suppressors, and potential novel drivers (i.e., genes detected as significant by Oncodrive3D and not annotated in the CGC or Fishy lists) (Supplementary Fig. 7c). Clumps in tumor suppressor genes are predominantly found in regions of the protein with low solvent accessibility, that is, buried within its hydrophobic core. In contrast, clumps identified in oncogenes follow a bimodal distribution, with a subset of clumps overlapping buried regions and another occurring at exposed areas, likely corresponding to binding pockets, interface surfaces, active sites, or similar functional regions. Also, clumps in tumor suppressors overlap the areas with the highest backbone rigidity (pLDDT), while clumps in oncogenes appear spread across a wider range of backbone rigidity. In agreement with this observation, mutations in clumps identified in tumor suppressor genes are predicted to have a higher destabilizing effect on protein folding (ΔΔG) compared to those in oncogenes. Finally, clumps in oncogenes tend to be more “pointy”, i.e., concentrating more mutations per residue involved in the clump, than those in tumor suppressor genes. Interestingly, clumps in potential novel driver genes exhibit properties that not only resemble those of oncogenes but are further amplified in certain aspects, such as greater solvent accessibility and lower stability change upon mutations, suggesting a possible gain-of-function mechanism underlying their role in cancer.

We were also interested in the analysis of the relationship of clump detection and the predicted alignment error (PAE) of pairs of residues involved in the clump. The underlying question was whether relatively high PAE in certain regions of the AlphaFold 2 models could result in spurious mutational 3D clusters identified by Oncodrive3D, just by virtue of their mutations being mistakenly considered part of the same volume. We had specifically designed an approach based on the PAE to define the probability that two residues became in contact in the structure (see Methods) with the aim to avoid this pitfall. For every significant gene, we computed a weighted average score of the PAE of all significant mutational 3D clusters, using the number of mutations observed in the corresponding volume as a weighting factor. Reassuringly, we observed that the majority of clumps of significant mutational 3D clusters identified across genes correspond to the lower end of the PAE distribution (Supplementary Fig. 7d).

### Supplementary Figures

### Supplementary Figure 1

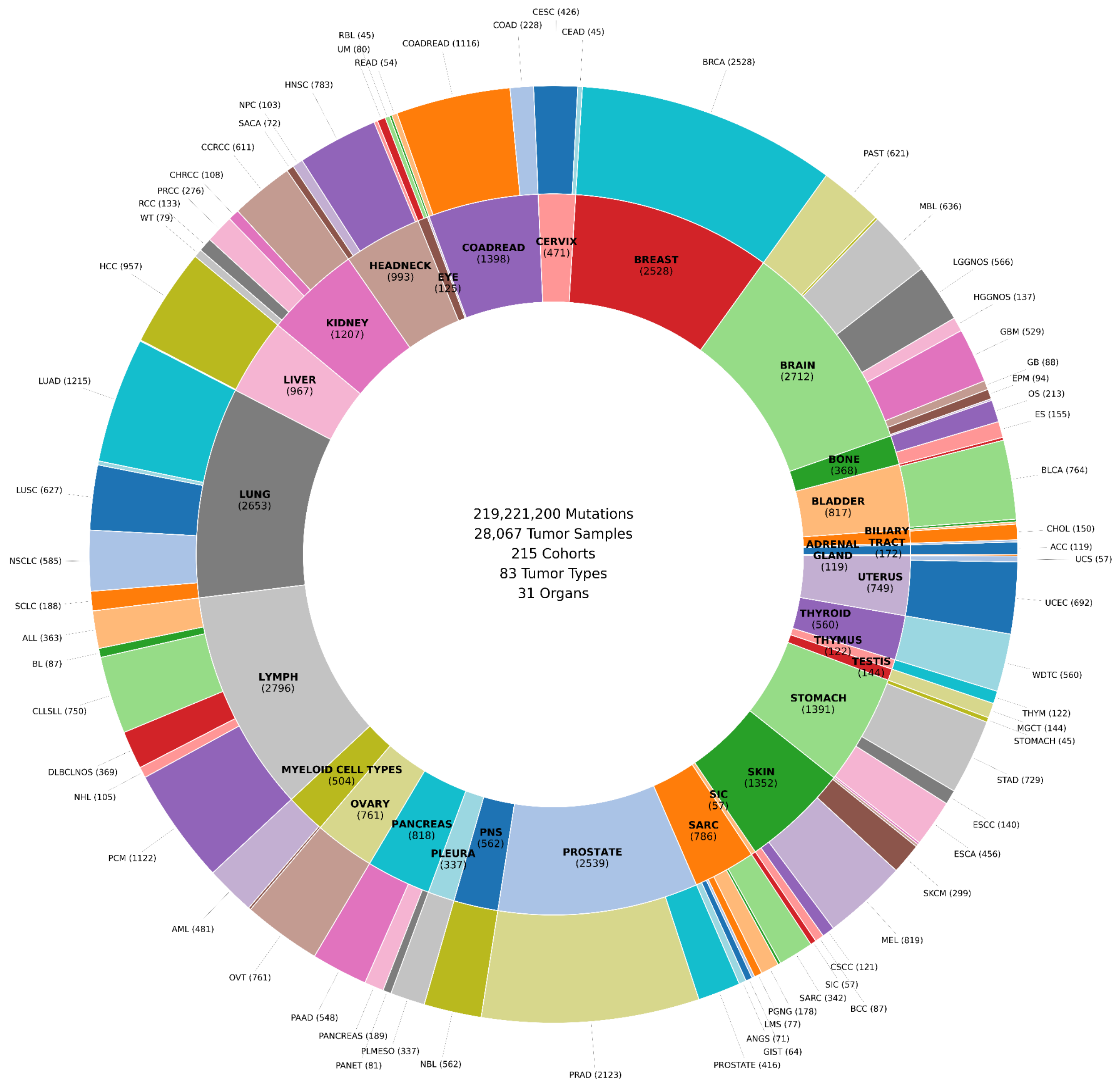

**Supplementary Figure 1. Cancer genomics datasets used to test the performance of Oncodrive3D**
The two concentric circles summarize the tumor types and number of samples corresponding to the 215 cohorts of tumors used to benchmark Oncodrive3D. The outer ring contains the names and number of samples analyzed for different tumor types, while the inner ring contains the names and number of samples analyzed for tumors of different organs. In parentheses, the number of samples processed for each cancer type or organ. Only labels for cancer types and organs with 40 samples or more are shown. A detailed description of the information in these datasets can be found in Supplementary Table 1. For acronyms of the tumor types, see intOGen (www.intogen.org) (3).

#### Supplementary Figure 2

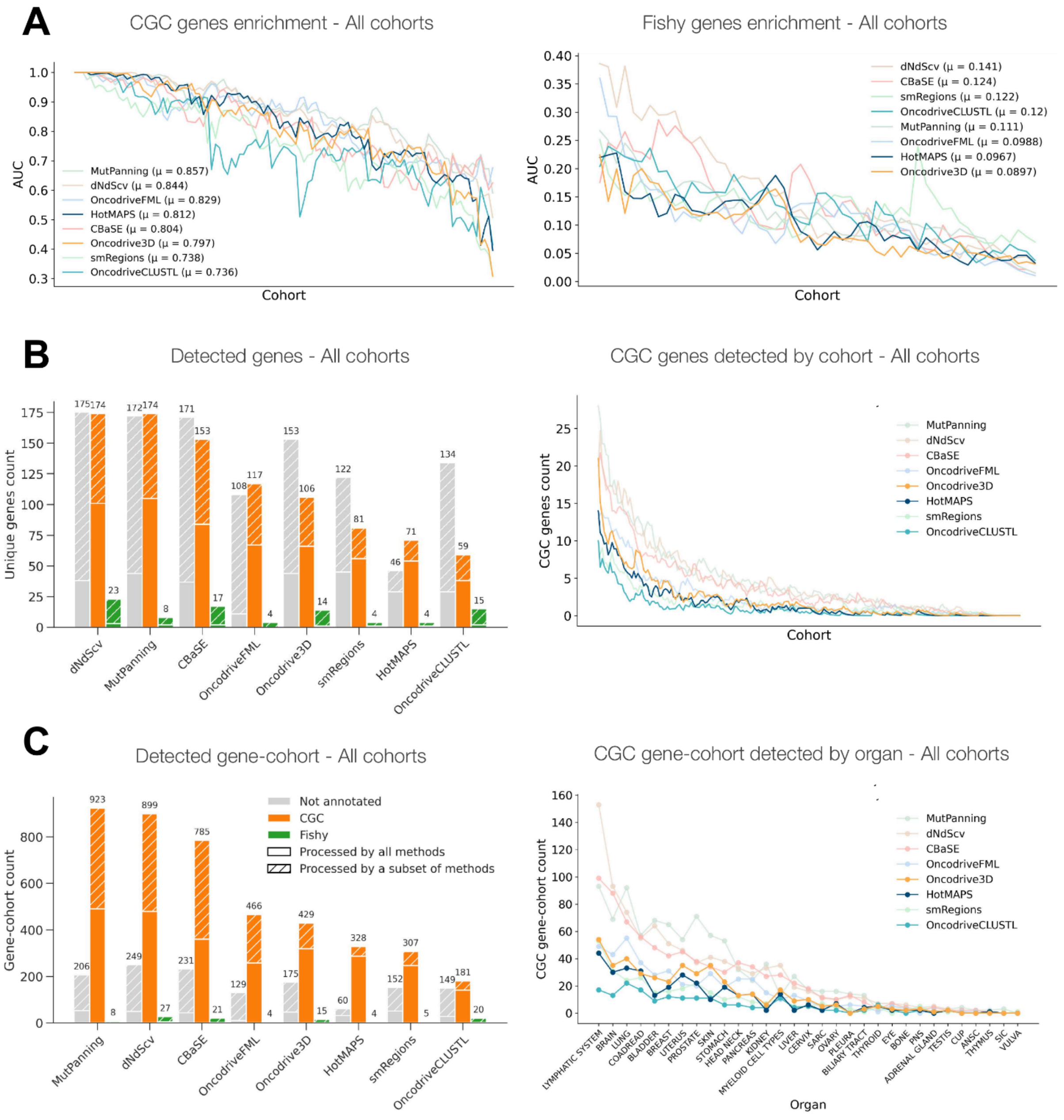

**Supplementary Figure 2. Benchmark of the sensitivity and specificity of Oncodrive3D**

A) AUC-CGC (left) and AUC-Fishy (right) of seven state-of-the-art driver discovery methods and Oncodrive3D across 215 intOGen cohorts. The mean AUC-CGC obtained by each method is shown in the graph.

B) Left, total number of genes that are either *bona fide* cancer drivers (CGC, orange), likely false positives (Fishy, green), or potential new drivers (Not annotated, gray) identified by seven state-of-the-art driver discovery methods and Oncodrive3D across 215 intOGen cohorts. Right, number of *bona fide* drivers identified by the seven state-of-the-art driver discovery methods and Oncodrive3D across 215 cohorts.

C) Left, total number of gene-cohort combinations involving genes that are either *bona fide* cancer drivers (CGC, orange), likely false positives (Fishy, green), or potential new drivers (Not annotated, gray) identified by seven state-of-the-art driver discovery methods and Oncodrive3D across 215 intOGen cohorts. Number of *bona fide* cancer gene-cohort combinations identified by the seven state-of-the-art driver discovery methods and Oncodrive3D across tumors of different organs.

In B and C, the solid segment of each bar corresponds to the fraction of genes that are analyzed (or processed) by all methods, while the segment with diagonal lines represents the fraction of genes analyzed only by a subset of the methods.

### Supplementary Figure 3

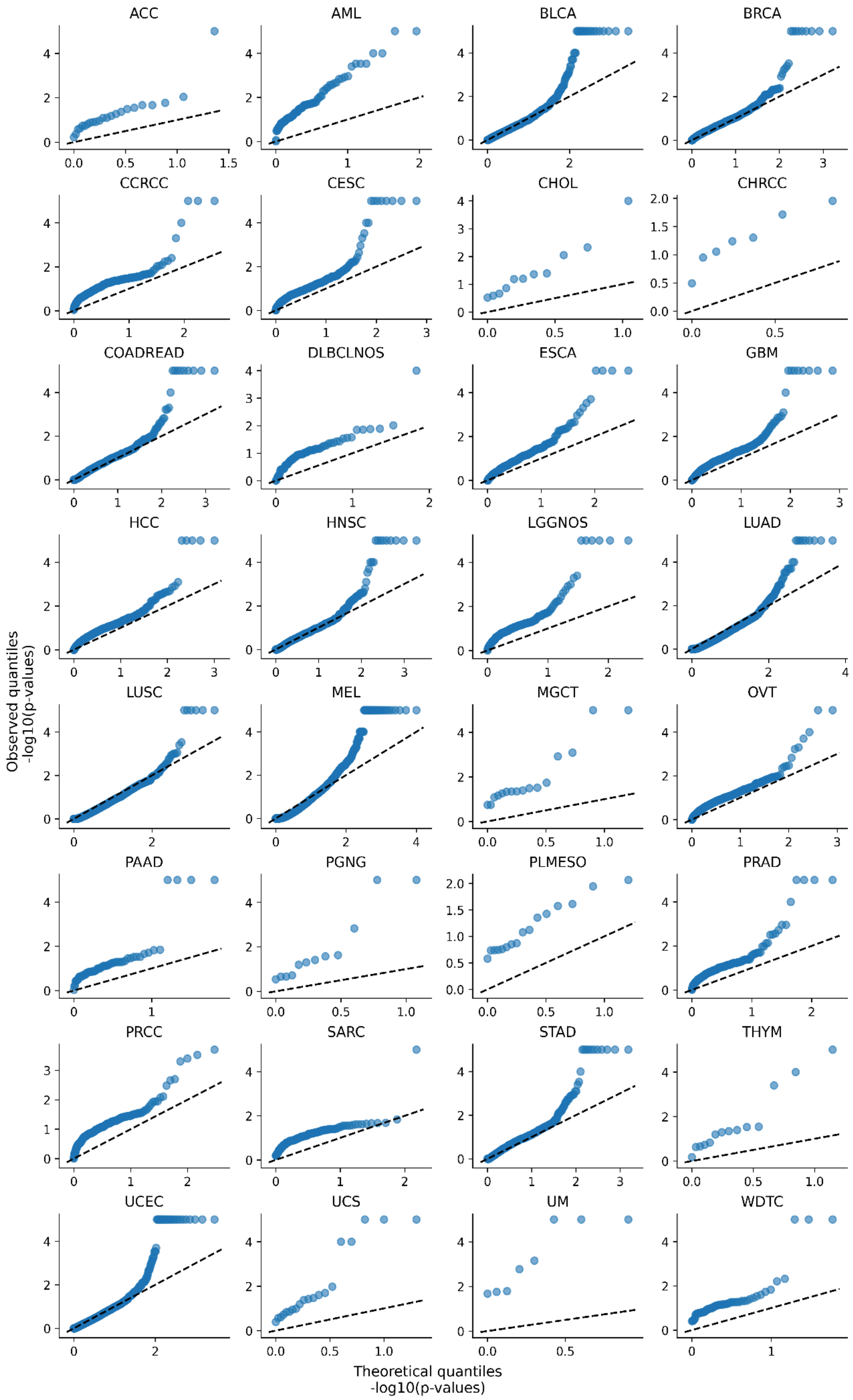

**Supplementary Figure 3. QQ-plots presenting observed and expected distribution of Oncodrive3D p-values in TCGA cohorts**

While in most cohorts the p-values of the test follow closely the expected uniform distribution (diagonal line), in some cohorts, in particular those where only few genes are tested, the distribution deviates from the expected. Importantly, no signs of inflation are observed in cohorts of tumors with high mutation burden, such as head and neck, lung adeno and lung squamous carcinomas, and melanomas.

#### Supplementary Figure 4

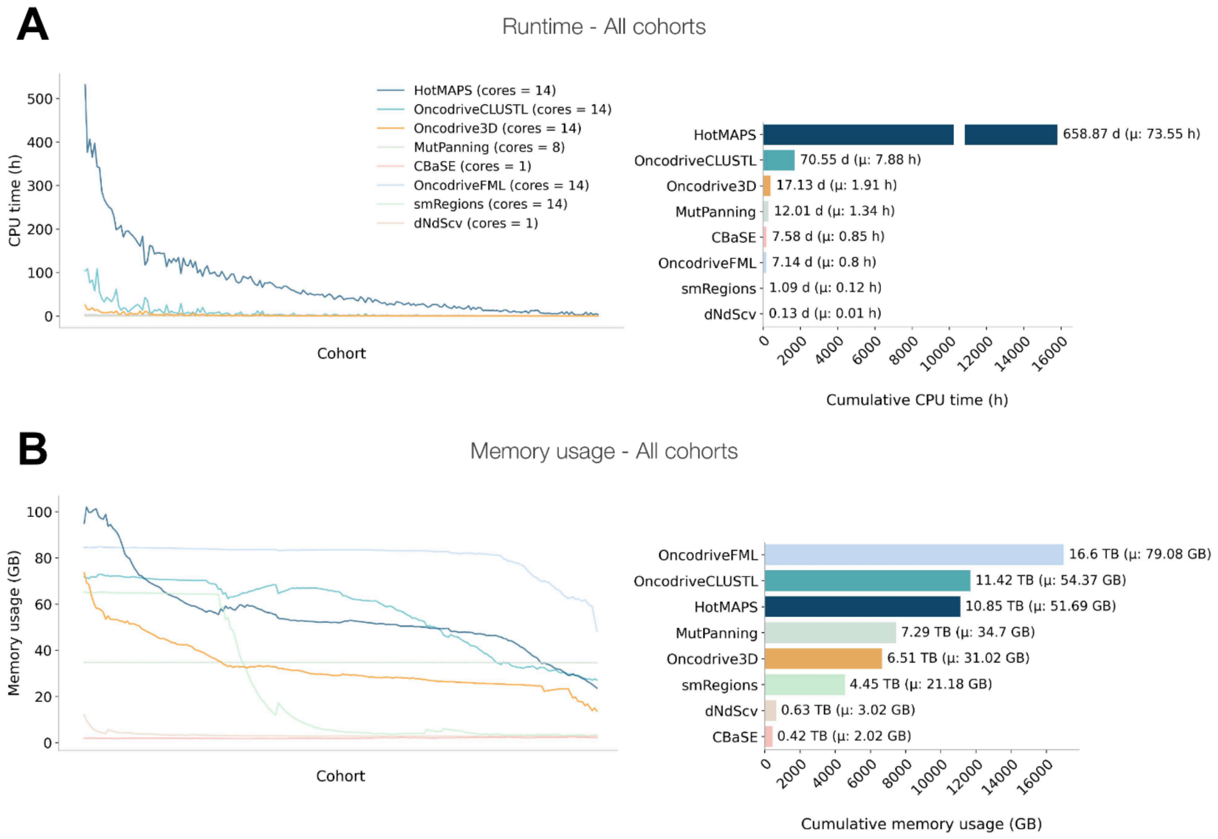

**Supplementary Figure 4. Benchmark of the computational efficiency of Oncodrive3D**

A) CPU-hours (left) or CPU-days occupied by seven state-of-the-art driver discovery methods and Oncodrive3D for the analysis of each (left) and all (right) 215 cohorts of tumors in intOGen.

B) Gigabytes (GB, left) or Terabytes (TB, right) of memory occupied by seven state-of-the-art driver discovery methods and Oncodrive3D for the analysis of each (left) and all (right) 215 cohorts of tumors in intOGen.

#### Supplementary Figure 5

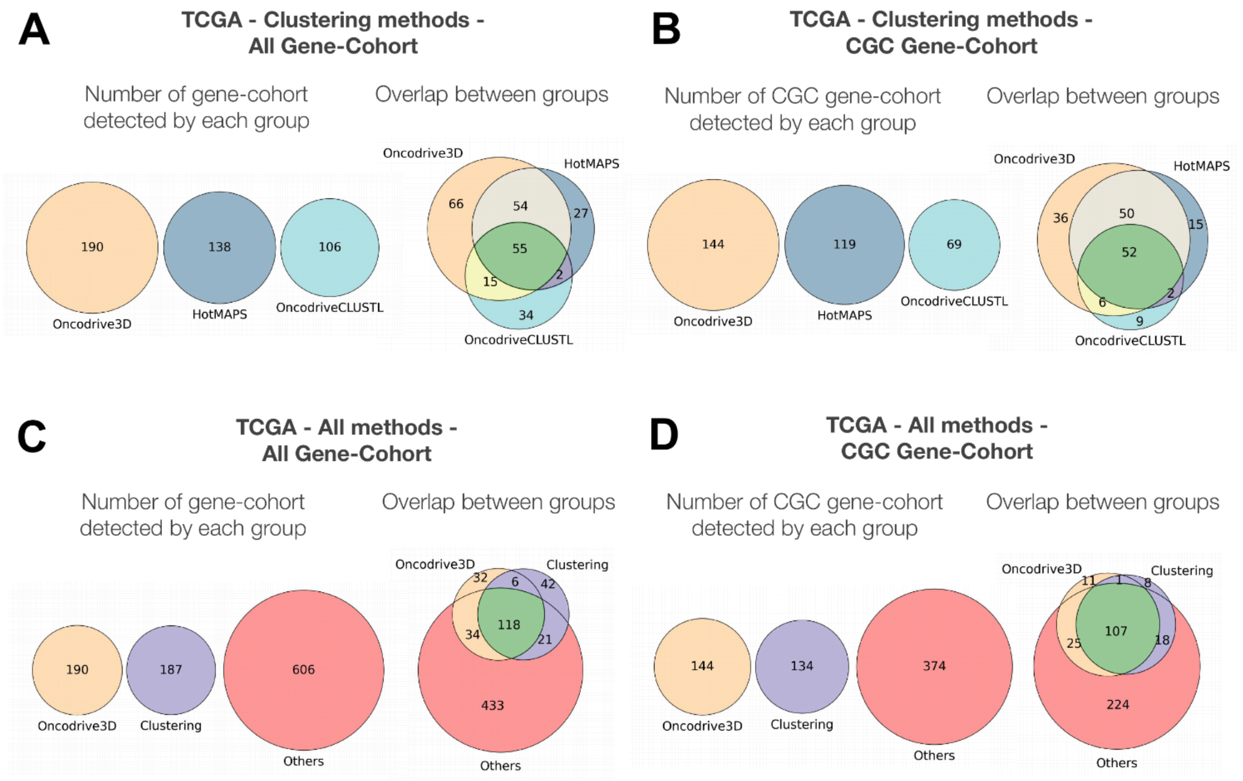

**Supplementary Figure 5. Further complementarity between Oncodrive3D and other driver discovery methods across 32 TCGA cohorts**

A,B) Overlap in the number of gene-cohort combinations involving any (A) or solely *bona fide* driver (annotated in the CGC, B) genes identified by Oncodrive3D, and two state-of-the-art driver discovery methods based on the identification of significant mutational clusters in the linear sequence (OncodriveCLUSTL) or the 3D protein structure (HotMAPS) across 32 TCGA cohorts.

C,D) Overlap in the number of gene-cohort combinations involving any (C) or solely *bona fide* driver (annotated in the CGC, D) genes identified by Oncodrive3D, and seven state-of-the-art driver discovery methods –grouped into two categories (Clustering and Others) based on the signals of positive selection of mutations that they exploit– across 32 TCGA cohorts.

#### Supplementary Figure 6

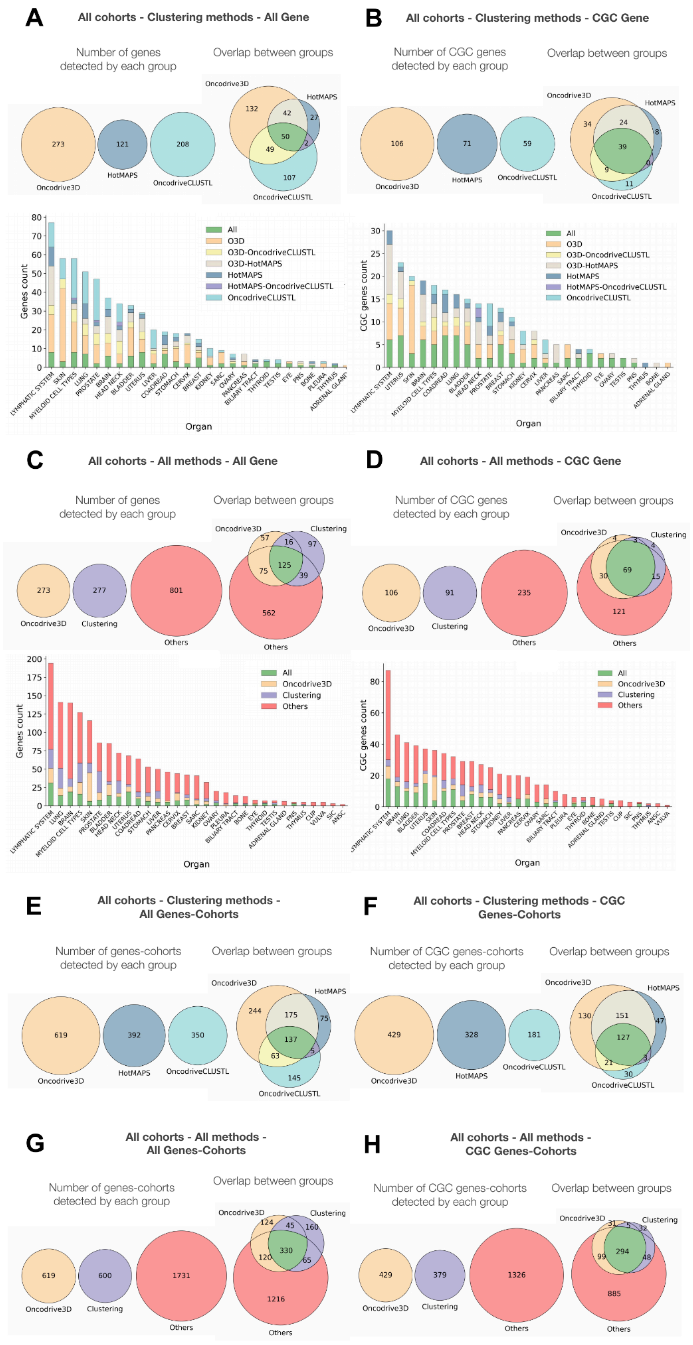

**Supplementary Figure 6. Complementarity between Oncodrive3D and other driver discovery methods across 215 intOGen cohorts**

A,B) Overlap in the total (A) or *bona fide* (annotated in the CGC, B) number of genes identified by Oncodrive3D, and two state-of-the-art driver discovery methods based on the identification of significant mutational clusters in the linear sequence (OncodriveCLUSTL) or the 3D protein structure (HotMAPS) across 215 intOGen cohorts. The intersections are represented as Venn diagrams (top panel) or bar plots (bottom panel).

C,D) Overlap in the total (C) or *bona fide* (annotated in the CGC, D) number of genes identified by Oncodrive3D, and seven state-of-the-art driver discovery methods –grouped into two categories (Clustering and Others) based on the signals of positive selection of mutations that they exploit– across 215 IntOGen cohorts. The intersections are represented as Venn diagrams (top panel) or bar plots (bottom panel).

E,F) Overlap in the number of gene-cohort combinations involving any (E) or solely *bona fide* driver (annotated in the CGC, F) genes identified by Oncodrive3D, and two state-of-the-art driver discovery methods based on the identification of significant mutational clusters in the linear sequence (OncodriveCLUSTL) or the 3D protein structure (HotMAPS) across 215 intOGen cohorts.

G,H) Overlap in the number of gene-cohort combinations involving any (G) or solely *bona fide* driver (annotated in the CGC, H) number identified by Oncodrive3D, and seven state-of-the-art driver discovery methods –grouped into two categories (Clustering and Others) based on the signals of positive selection of mutations that they exploit– across 215 intOGen cohorts.

### Supplementary Figure 7

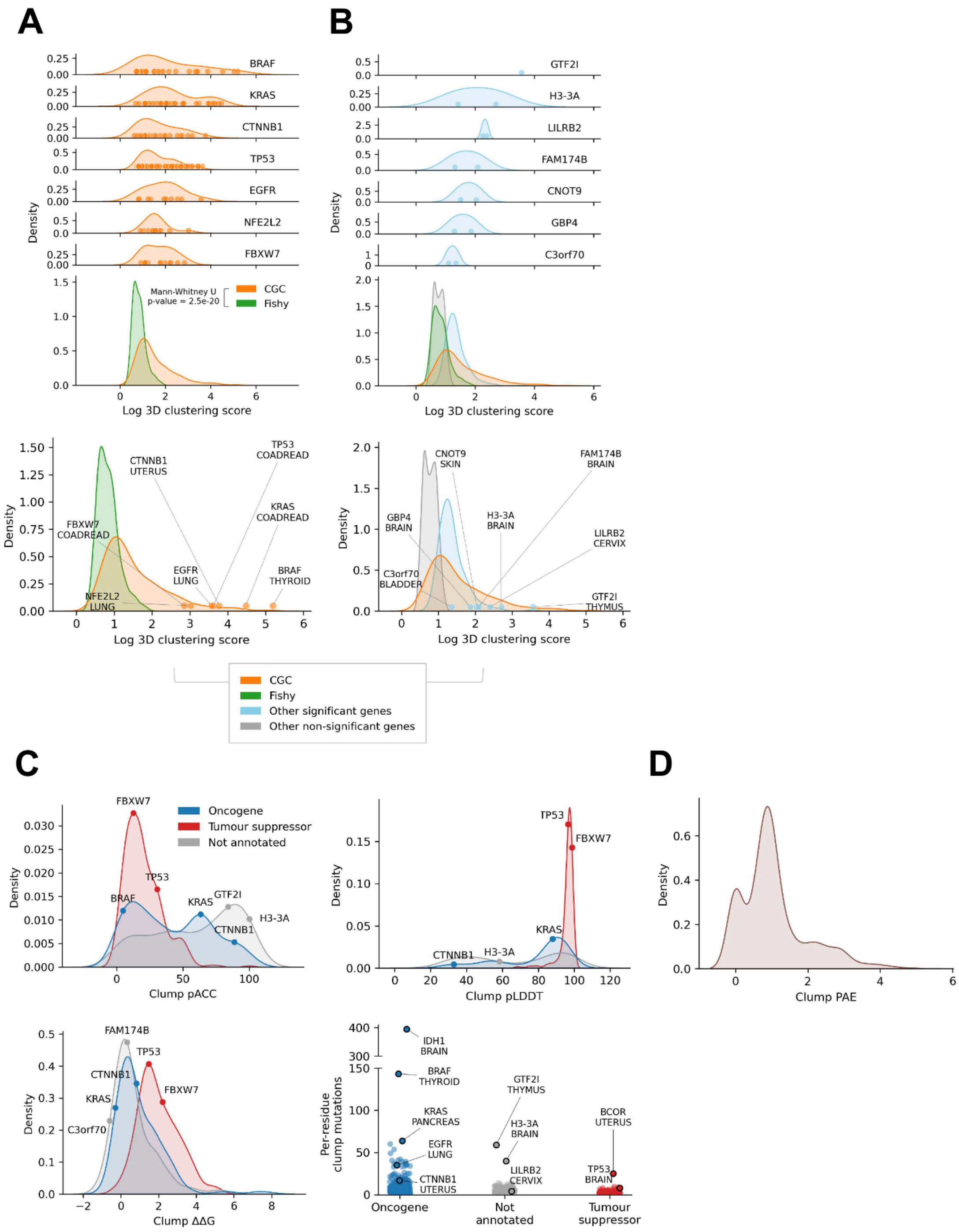

**Supplementary Figure 7. Characteristics of genes with significant 3D mutational clusters**

A) Distribution of 3D clustering scores for all significant residues identified by Oncodrive3D in seven selected *bona fide* cancer genes (top). The middle panel shows the distribution of 3D clustering scores, defined for each gene as the score of its most significant residue, across all CGC and Fishy genes in 215 intOGen cohorts. The bottom panel is a magnified version of the middle plot, annotated with seven selected *bona fide* driver gene-cohort combinations within the distribution. For each *bona fide* cancer driver, the gene-cohort combination where that gene achieved the highest clustering score is highlighted.

B) Same as A, but focused on potentially novel driver genes (i.e., not annotated in the CGC and outside the Fishy list) identified by Oncodrive3D. As in A, the top plot presents seven exemplary genes, while in the middle and bottom plots, the distribution of 3D clustering scores of all novel driver genes and all not significant genes are added. Importantly, as shown in the middle plot, while the distribution of 3D clustering scores of residues of non significant genes overlaps with that corresponding to Fishy genes, the distribution of potential novel drivers (“Other significant genes” in the plot) is clearly distinct and overlaps that of CGC genes. For clarity, the distribution of Fishy genes is removed from the bottom plot.

C) Distribution of solvent accessibility (top left), rigidity of the backbone (top right), ΔΔG resulting from amino acid substitution in clusters (bottom left), and number of observed mutations per residue in the cluster (bottom right) across clumps of significant 3D mutational clusters in oncogenes, tumor suppressor, and unclassified genes (novel candidate driver genes).

D) Distribution of average AlphaFold predicted aligned error (PAE) (right) for all clumps of significant mutational 3D clusters identified by Oncodrive3D across the 215 intOGen cohorts.

#### Supplementary Figure 8

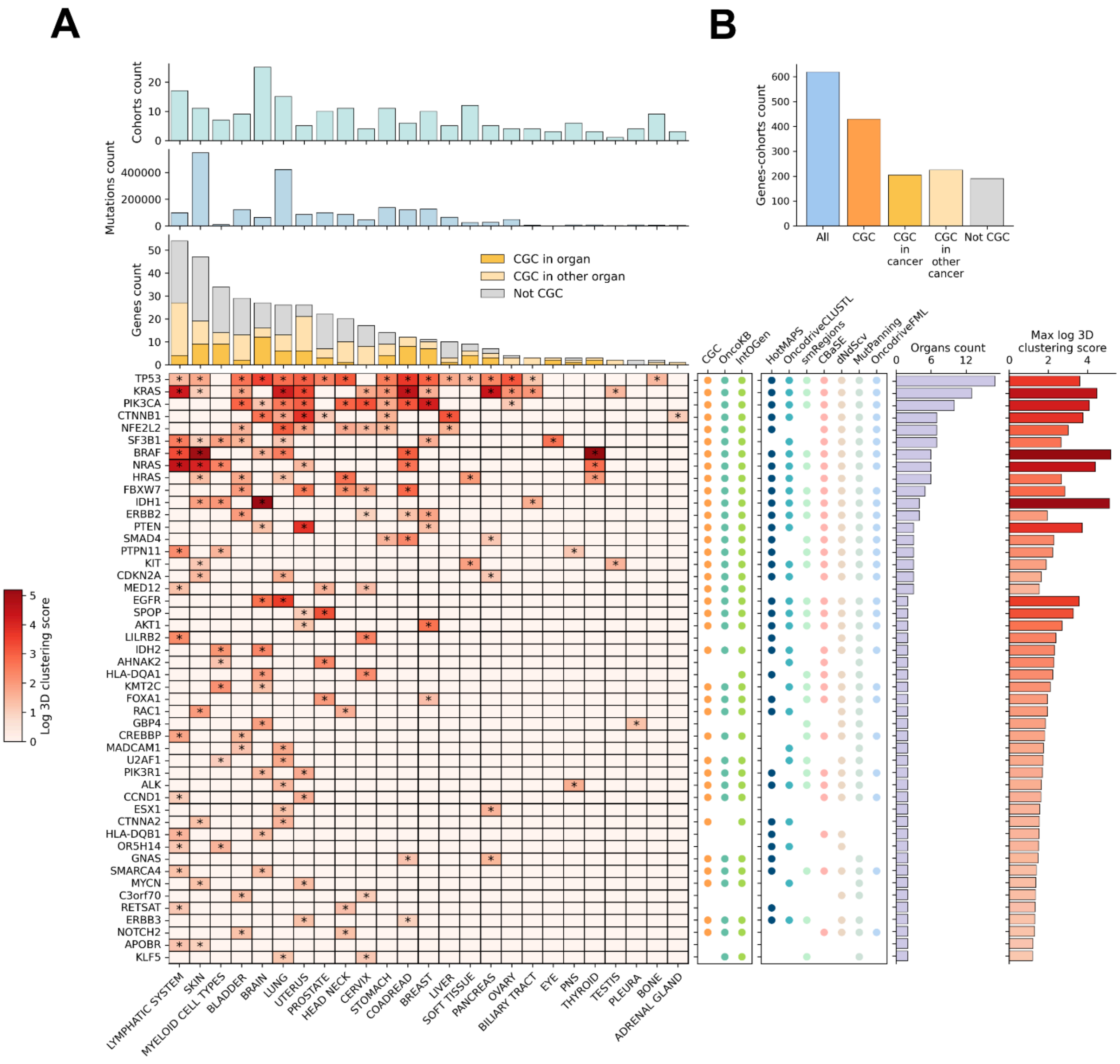

**Supplementary Figure 8. Top genes identified by Oncodrive3D across 215 intOGen cohorts**
A) The central heatmap presents the top significant 48 genes (in terms of number of tumors from different organs where they are identified by Oncodrive3D; cells with an asterisk). The color of the cells represent the 3D clustering score of the residue with the lowest p-value in each gene. The two bar plots above the heatmap represent the total number of missense mutations in each cohort and the total number of genes (annotated in the CGC for the tumor type of the cohort in question, for other tumor types, or not annotated in the CGC) identified as bearing significant 3D mutational clusters by Oncodrive3D. The first rectangular panel by the right side of the heatmap denote whether the gene is annotated in any of two catalogues of *bona fide* cancer genes (CGC, OncoKB) (4, 5), or has been identified by the intOGen pipeline (3). The second rectangular panel denotes which of the driver discovery methods in the intOGen pipeline has identified the gene as a potential cancer driver. The bars on the right represent the number of organs in which each gene has been found to bear significant mutational 3D clusters by Oncodrive3D (Organ count) and the maximum 3D clustering score, displayed on a log scale, that each gene has achieved across all cohorts (Max log 3D clustering score).
B) Total number of gene-cohort combinations identified as bearing significant 3D clusters of mutations by Oncodrive3D.
In the heatmap only 27 organs (those for which at least one driver gene is identified by Oncodrive3D) are included.

### Supplementary Figure 9

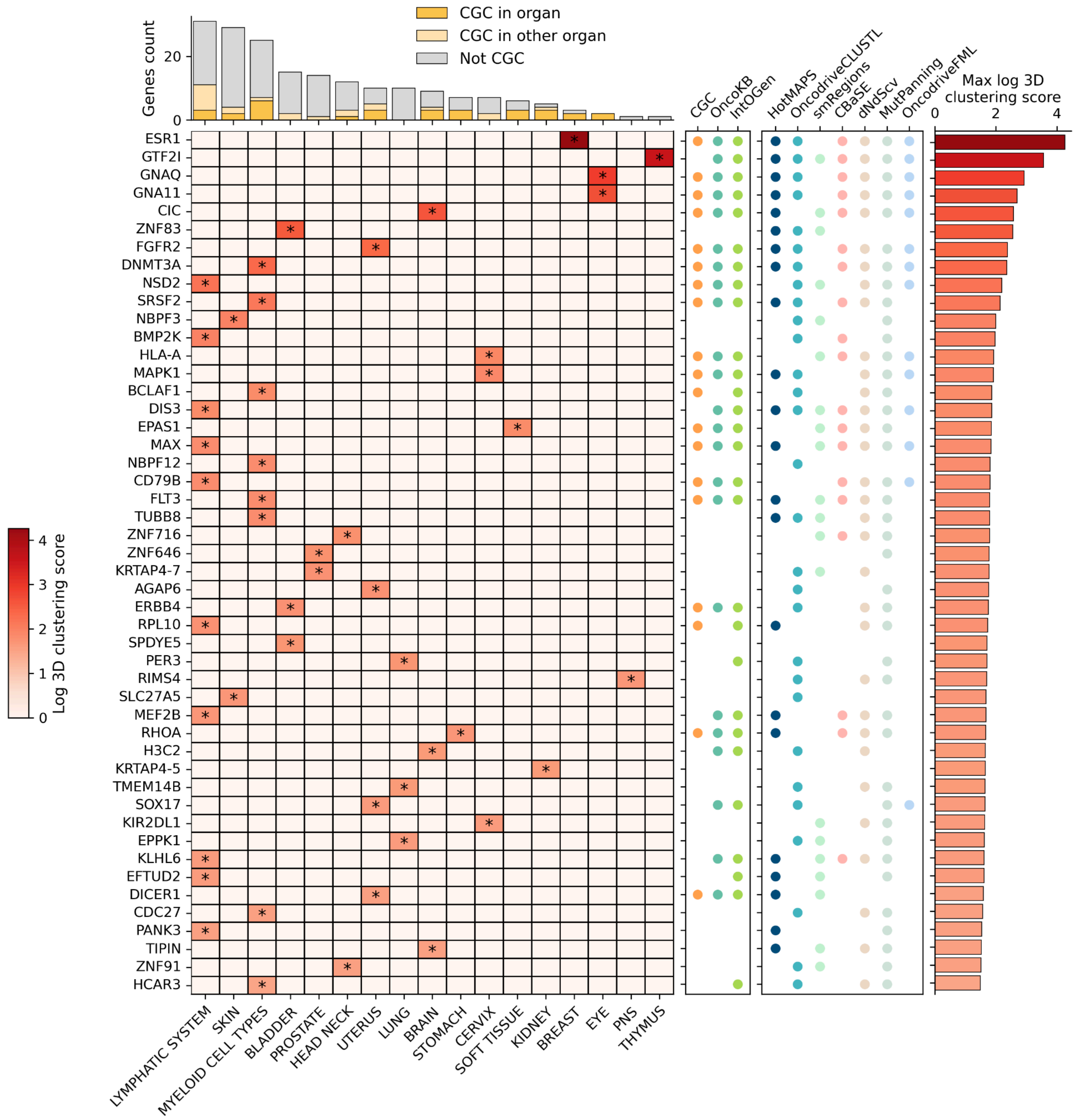

**Supplementary Figure 9. Selected genes with significant mutational 3D clusters in only one cohort**
The legend is as in Supplementary Figure 8a but this Figure includes only genes that have been detected as significant by Oncodrive3D in only one cohort.

#### Supplementary Figure 10

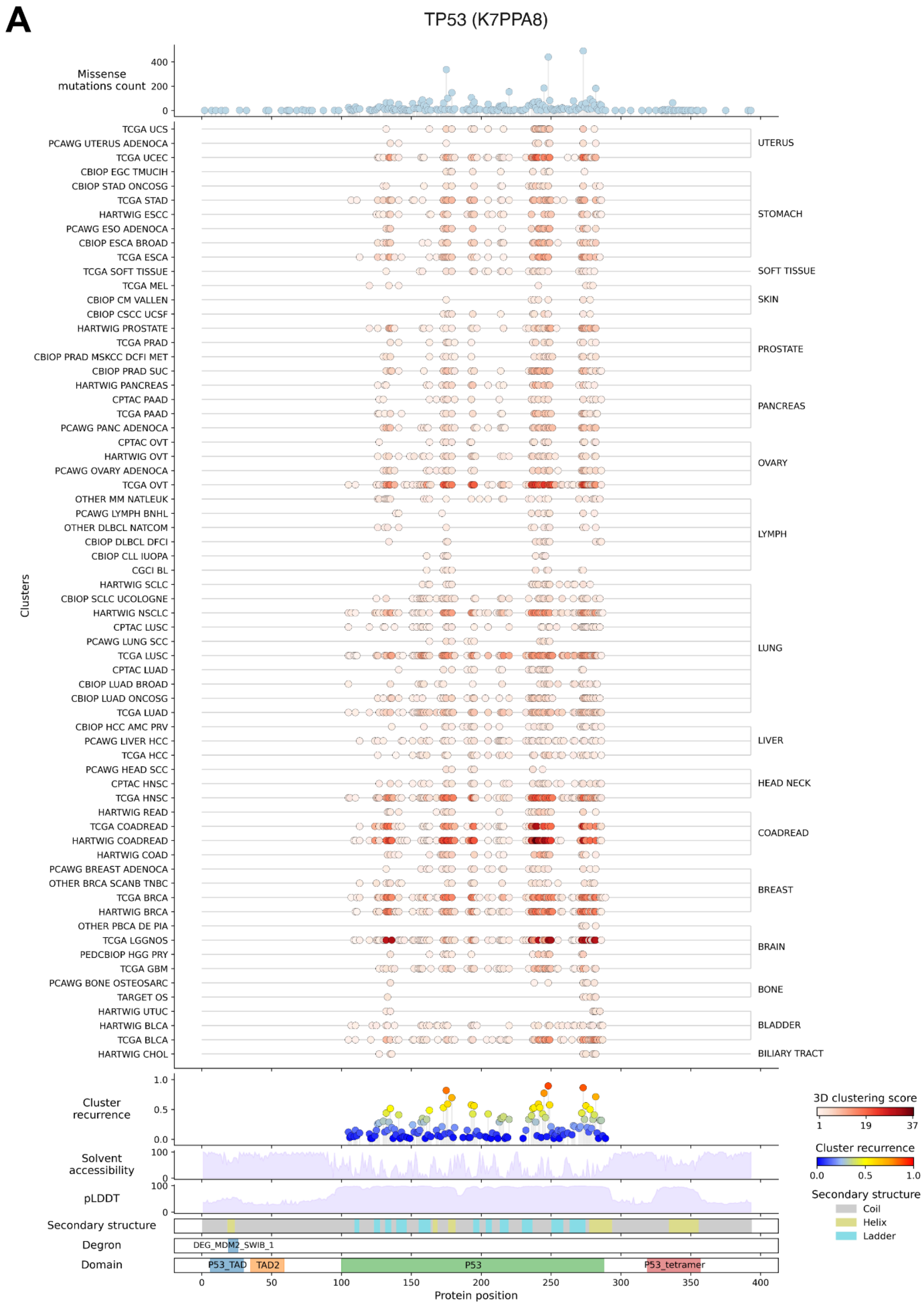

B

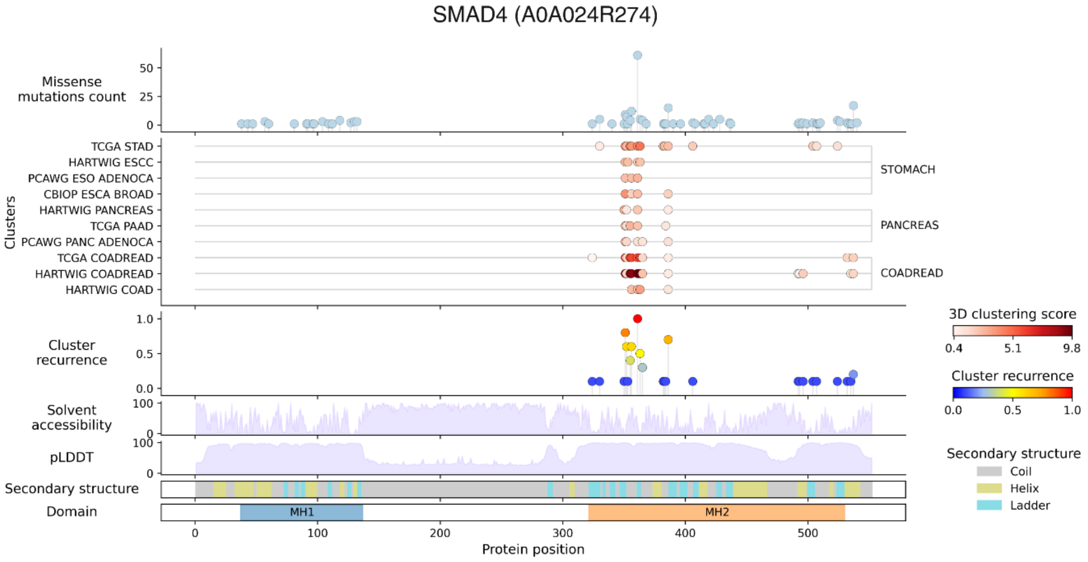
SMAD4 (A0A024R274)
Missense mutations count
Clusters
TCGA STAD
HARTWIG ESCC
PCAWG ESO ADENOCA
CBIOP ESCA BROAD
HARTWIG PANCREAS
TCGA PAAD
PCAWG PANC ADENOCA
TCGA COADREAD
HARTWIG COADREAD
HARTWIG COAD
STOMACH
PANCREAS
COADREAD
Cluster recurrence
Solvent accessibility
pLDDT
Secondary structure
Domain
MH1
MH2
Protein position
3D clustering score
0.4
5.1
9.8
Cluster recurrence
0.0
0.5
1.0
Secondary structure
Coil
Helix
Ladder

C

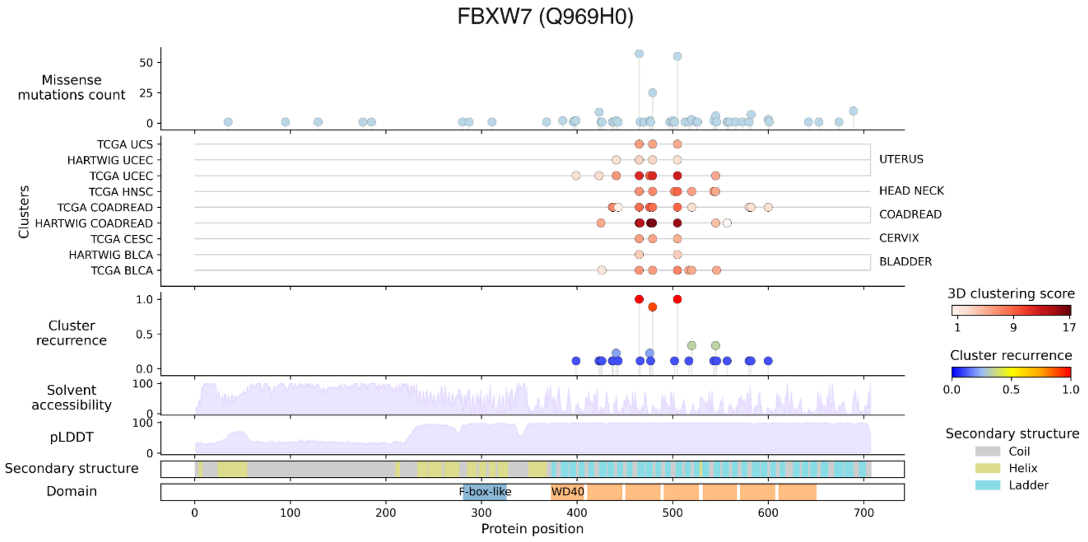
FBXW7 (Q969H0)
Missense mutations count
Clusters
TCGA UCS
HARTWIG UCEC
TCGA UCEC
TCGA HNSC
TCGA COADREAD
HARTWIG COADREAD
TCGA CESC
HARTWIG BLCA
TCGA BLCA
UTERUS
HEAD NECK
COADREAD
CERVIX
BLADDER
Cluster recurrence
Solvent accessibility
pLDDT
Secondary structure
Domain
F-box-like
WD40
Protein position
3D clustering score
1
9
17
Cluster recurrence
0.0
0.5
1.0
Secondary structure
Coil
Helix
Ladder

**Supplementary Figure 10. Recurrence of residues with significant mutational 3D clusters in *bona fide* cancer genes**

Recurrence of residues with clusters identified in TP53 (A) and SMAD4 (B) and FBXW7 (C) across cohorts of tumors affecting different organs.

In the three graphs, the top panel presents the number of mutations affecting each residue of the protein. The second panel presents the residues identified as bearing significant mutational 3D clusters of mutations by Oncodrive3D in each cohort of tumors analyzed, with the color representing the 3D clustering score of each of them. The third panel represents the recurrence of each cluster (fraction of cohorts analyzed where the cluster is significant). The 4 panels below present annotations of the protein structure that support the interpretation of the functional relevance of the clusters; from top to bottom: solvent accessibility (6, 7), backbone rigidity (6, 7), secondary structure (https://github.com/realbigws/PDB_Tool) and functional domains (8).

### Supplementary Figure 11

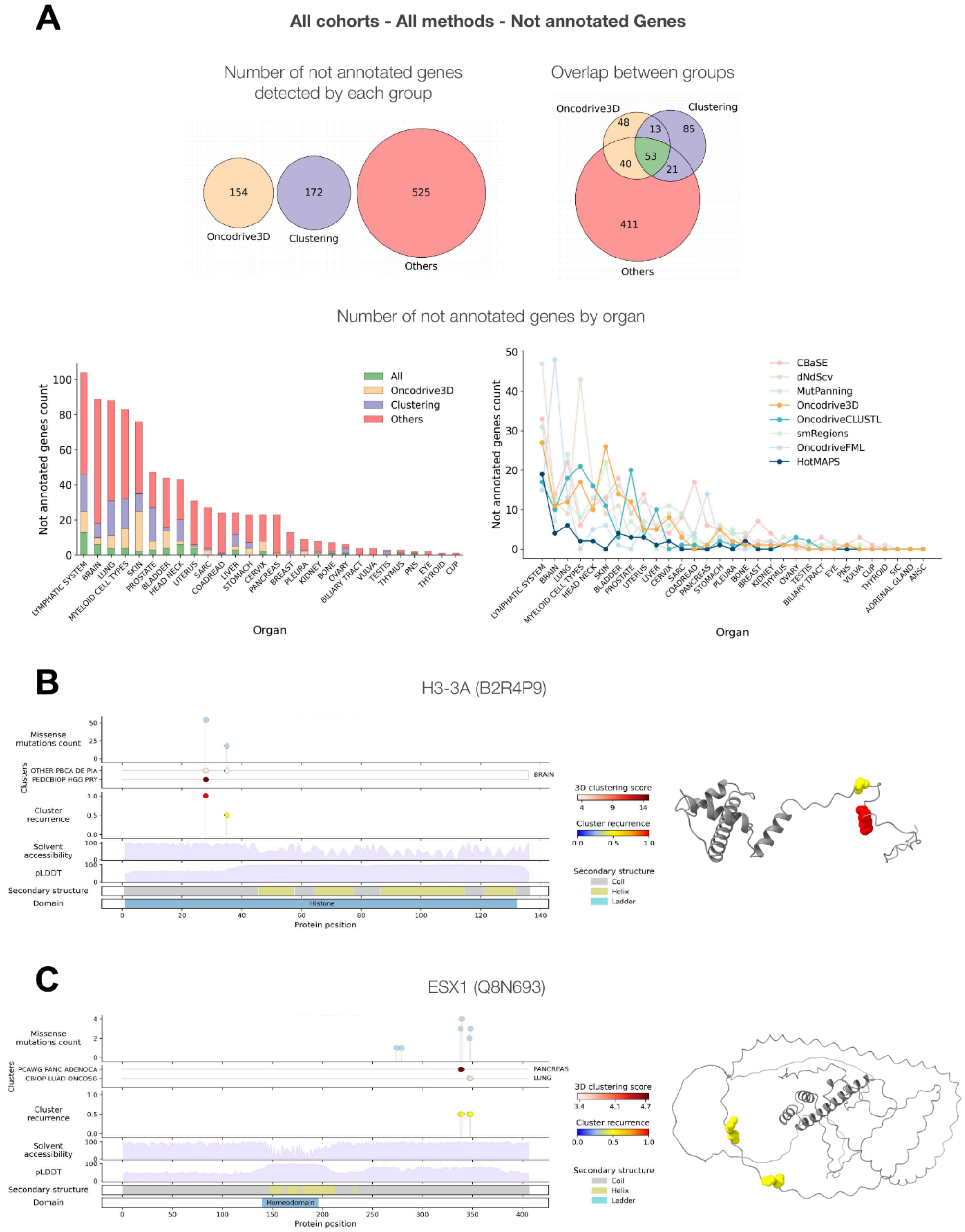

**Supplementary Figure 11. Significant mutational 3D clusters in three potential novel cancer genes**
A) We focused on 53 potentially novel cancer driver genes that were identified as significant by Oncodrive3D, at least one other clustering-based method and another method based on a different signal of positive selection. The legend of this panel is analogous to that in Supplementary Figure 6a.
B) Conservation of 3D mutational clusters identified in H3-3A (B) and ESX1 (C) across cohorts of tumors affecting different organs. These two panels are analogous to those in Supplementary Figure 10a,b and c.

### Supplementary Figure 12

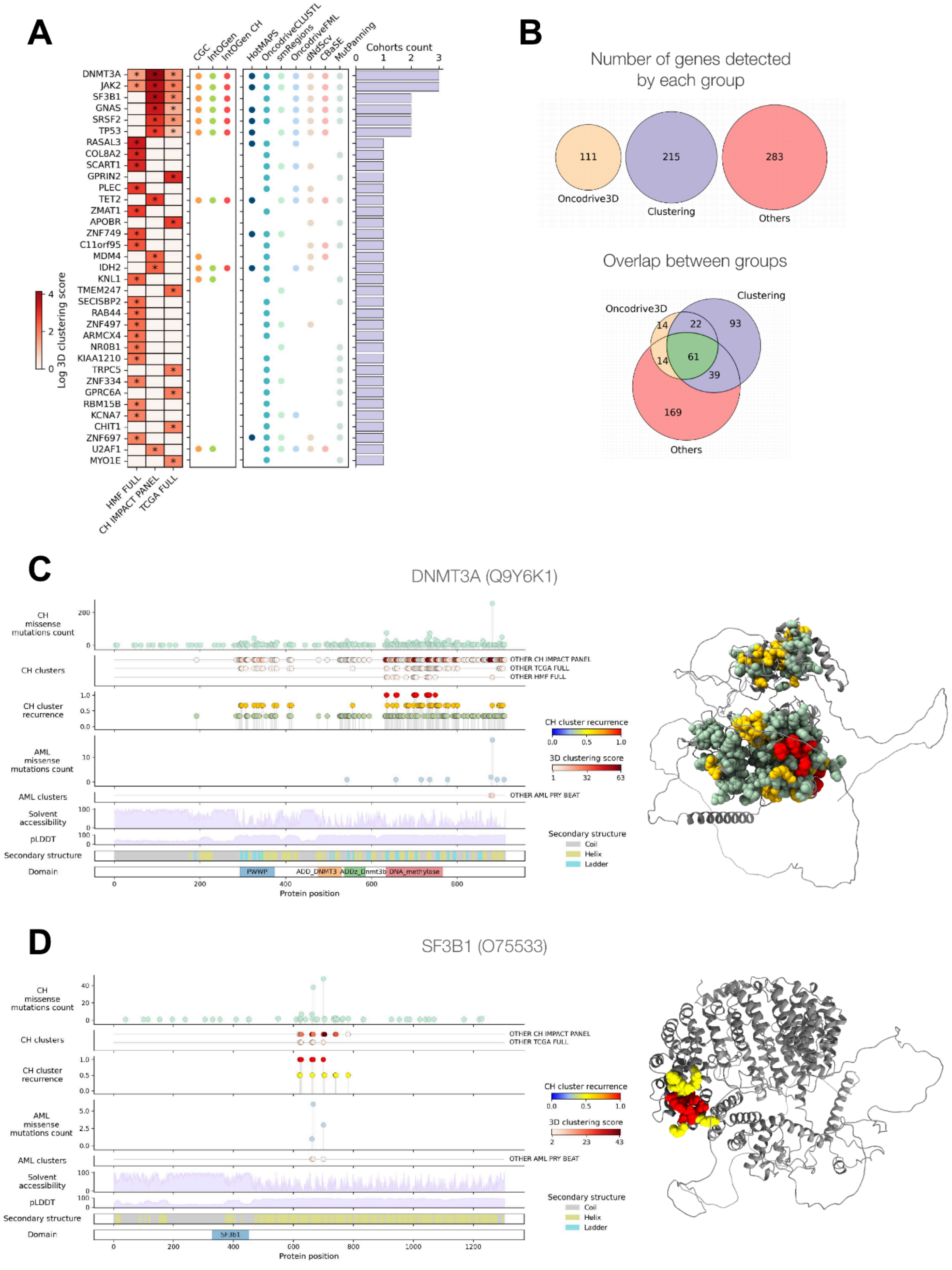

**Supplementary Figure 12. Oncodrive3D identifies significant 3D clusters of mutations in CH drivers**

A) Result of the application of Oncodrive3D to three datasets of blood somatic mutations identified in the same number of cohorts through reverse mutation calling (9). This logic of this panel is analogous to that presented in Supplementary Figure 8a.

B) Complementarity in the identification of CH driver genes of driver discovery methods based on different signals of positive selection.

C,D) Recurrence of 3D mutational clusters identified in DMT3A (C) and SF3B1 (D) across 3 cohorts of donors and cohorts of AML.

### Supplementary Tables description

**Supplementary Table 1**: Description of cohorts of tumors and CH used to test Oncodrive3D
**Supplementary Table 2**: List of likely false positive “Fishy” genes used to evaluate the specificity of driver discovery methods
**Supplementary Table 3**: Significant genes identified by Oncodrive3D across 215 cohorts of tumors and 3 cohorts of CH
